# Regulation of *Sox8* through lncRNA *Mrhl* mediated chromatin looping in mouse spermatogonia

**DOI:** 10.1101/2021.10.05.463295

**Authors:** Bhavana Kayyar, Anjhana C. Ravikkumar, Utsa Bhaduri, M.R.S Rao

## Abstract

Sox8 is a developmentally important transcription factor that plays an important role in sex maintenance and fertility of adult mice. In the B-type spermatogonial cells, *Sox8* is regulated by the lncRNA *Mrhl* in a p68-dependant manner under the control of the Wnt signalling pathway. The downregulation of *Mrhl* leads to the meiotic commitment of the spermatogonial cells in a *Sox8-*dependant manner. While the molecular players involved in the regulation of transcription at the *Sox8* promoter have been worked out, our current study points to the involvement of the architectural proteins CTCF and cohesin in mediating a chromatin loop that brings the *Sox8* promoter in contact with a silencer element present within the gene body in the presence of lncRNA *Mrhl* concomitant with transcriptional repression. Further, lncRNA *Mrhl* interacts with the *Sox8* locus through the formation of a DNA:DNA:RNA triplex which is necessary for the recruitment of PRC2 to the locus. The downregulation of lncRNA *Mrhl* results in the promoter-silencer loop giving way to a promoter-enhancer loop. This active transcription associated chromatin loop is mediated by YY1 and brings the promoter in contact with the enhancer present downstream of the gene.

## Introduction

*Sox8* is a developmentally important transcription factor that is critical for the maintenance of adult male fertility. *Sox8* knockout mice become progressively infertile because of age-related degeneration of spermatogenesis (O’Bryan et al, 2008). The Sertoli specific deletion of *Sox9* (another essential transcription factor involved in mammalian sex determination) in *Sox8* null embryonic mice results in failure to achieve the first wave of spermatogenesis (Barrionuevo et al., 2010). The deletion of both *Sox8* and *Sox9* in adult sertoli cells results in testis to ovarian genetic reprogramming (Barrionuevo et al, 2016) while *Sox8* alone is sufficient for ovarian to testicular genetic reprogramming in the absence of R-spondinl (Richardson et al, 2020)

Regulation of gene expression in mammals is a complex interplay of intricately controlled events. While decades of work has brought about some understanding of these regulatory events including the role of transcriptional factors, the role of proximal and distal regulatory elements, the long non-coding RNAs are the latest entrants to this hotbed of research. It is emerging that lncRNAs are inextricably involved in every step of gene regulation.

*Mrhl* lncRNA is a 2.4kb long mono-exonic transcript that is transcribed from an independent transcription unit within the 15th intron of the phkb gene in mice (Nishant et al, 2004). This chromatin-bound lncRNA regulates the expression of multiple genes in the mouse spermatogonial cells in a Wnt signalling dependent manner. *Sox8* is one of the three genes that the lncRNA regulates by binding to their promoters in a p68-dependent manner (Akhade et al, 2014).

Our group has previously deciphered the gene regulatory mechanism in play at the promoter of *Sox8* in mouse spermatogonial cells. *Mrhl* binds at the *Sox8* promoter ∼140bp upstream of the Transcription start Site (TSS) and the Mad-Max transcription factors along with the co-repressors Sin3a and HDAC1 are also bound at the *Sox8* promoter close to the *Mrhl* binding site in the presence of *Mrhl*. Wnt signalling-mediated transcriptional regulation of *Sox8* involves substantial changes in the chromatin dynamics of the promoter. There is a concomitant transcriptional activation of *Sox8* expression with downregulation of *Mrhl* and associated changes include the Mad-Max complex being replaced by the Myc-Max transcription factors and increased levels of H3K4me3 and H3K9ac histone modifications and decreased levels of H3K27me3 modification at this locus. Simultaneously, beta-catenin binds at the Wnt responsive element present at the promoter (Kataruka, S. et al., 2017). Activation of the Wnt signalling cascade in Gcl-spg cells results in their meiotic commitment marked by the increase in the levels of pre-meiotic and meiotic markers (Akhade, V.S. et al., 2016) including that of the master regulator, Stra8, in a *Sox8*-dependent manner.

While chromatin associated lncRNAs interact with DNA through protein bridges, an alternative mode of interaction is by directly binding to DNA. The potential of lncRNAs to form hybrid structures such as DNA:DNA:RNA triplexes to directly interact with DNA have been explored in the past decade. LncRNAs including *Meg3, KHPS1* and *PARTICLE* have been reported to form triplex structures at genomic regions having AG rich motifs (Mondal et al, 2015, Postepska-Igielska et al, 2015, O’Leary et al, 2015). In addition to acting as tethers, triplexes have been shown to act as platforms for the recruitment of DNA methyltransferase complex DNMT3b by pRNA at the *rDNA* locus or the polycomb repressive complex PRC2 by lncRNAs *MEG3* and *Fendrr* (Schmitz et al, 2010, Mondal et al, 2015, Grote and Herrmann, 2013) and thereby influence the expression of genes in the vicinity.

Recent advances indicate a role for lncRNAs in bringing together distant gene regulatory elements in 3D space to regulate gene expression. CTCF, the master architectural protein in mammalian cells, mediates interactions between distant genomic elements, resulting in context-dependent functional outcomes. The cohesin complex is essential in stabilising these CTCF mediated 3D contacts while various proteins such as Ying-Yangl (YY1), DDX5/p68 and CHD8 among others supplement CTCF’s role in genome organization (Ong and Corces, 2014). CTCF can bind to lncRNAs through its RNA-binding domain (Saldaña-Meyer et al, 2014) and the deletion of the RNA-binding domain within CTCF compromises its ability to homodimerise - an essential ability to enable chromatin looping (Saldaña-Meyer et al, 2019). The lncRNA *CCAT1-L* associates with CTCF to bring the MYC promoter in contact with the enhancer 515Kb away from where it is transcribed itself (Xiang et al, 2014). The role of the CTCF-cohesin complex along with lncRNA *SRA* and the DEAD-box RNA helicase DDX5/p68 in maintaining imprinting at the *H19/Igf2* has been extensively studied (Yao et al, 2010).

In the present study, we show that *Mrhl*, in association with the nuclear organizing factors CTCF, cohesin and YY1, mediates the formation of dynamic chromatin loops at the locus. In addition, *Mrhl* forms a triplex at the *Sox8* promoter and contributes to the regulation of gene expression through the triplex mediated recruitment of PRC2.

## Results

### *Mrhl* forms a triplex at the *Sox8* locus

*Sox8* is transcribed from a bidirectional promoter on chromosome 17. The binding of *Mrhl* at the promoter of *Sox8*, 140bp upstream of the TSS, is dependent on DDX5/p68 (Fig 1A). Since multiple chromatin associated lncRNAs interact with DNA through the formation of DNA:DNA:RNA triple helix structures, it was of interest to investigate if *Mrhl* lncRNA also interacts with the *Sox8* locus through the formation of triplex.

**Fig 1:**
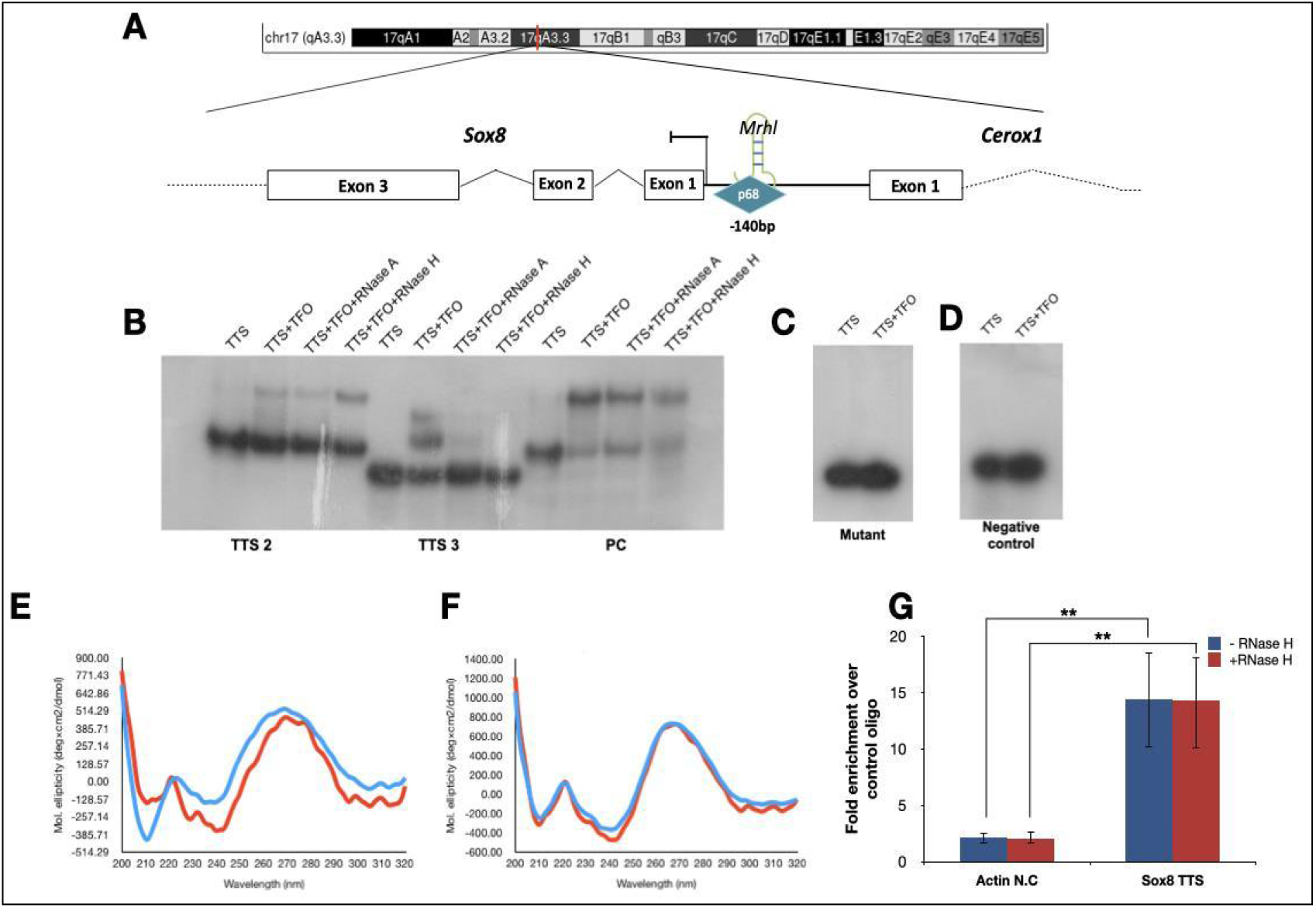
Mrhl interacts at the Sox8 locus. **(A)** The mouse Sox8 locus on chromosome 17 where Mrhl lncRNA binds at the bidirectional promoter 140bp upstream of the TSS in a p68-dependent manner to maintain Sox8 in the transcriptionally repressed state. **(B)** EMSA performed for the indicated DNA oligo incubated with RNA oligo TFO1. The lanes have the reaction components as indicated above them **(C)** EMSA performed with mutant TTS2 and RNA TFO1 where no shift in mobility is observed **(D)** EMSA for negative control TFO/TTS pair **(E)** Artificial spectrum generated by summation of individual CD spectra recorded for TTS2 only and NC TFO only (red) overlaid with CD spectrum of triplex reaction for the oligonucleotides, **(F)** Artificial spectrum generated by summation of individual CD spectra recorded for TTS2 only and TFO1 only (red) overlaid with CD spectrum of triplex reaction for the same oligonucleotides. Plots are an average of 4 independently recorded spectra. **(G)** Results of the in-nucleus triplex pulldown assay show significant enrichment of the Sox8 ITS region in the TFO1 oligo associated chromatin fraction over NC TFO associated chromatin fraction both without and with RNaseH digestion. Data in the graph has been plotted as mean ±S.D., N=3. *** P ≤ 0.0005, ** P ≤ 0.005, * P ≤ 0.05, N.S - Not significant (two-tailed Student’s t test)

Majority of the lncRNA:DNA triplexes have been reported to form at AG- rich regions. The region ∼1.2kb upstream of the *Sox8* TSS harbours ∼50 AG-repeats and is a prime candidate for triplex formation. Initial *in silico* predictive analysis was done using the Triplexator tool. Triplex Forming Oligo (TFO)-Triplex Target Sites (TTS) pairs with a score of 10 and above (Table 1) were considered for further experimentation. The TTS sequences were found to be overlapping within a 33bp DNA sequence between the regions -1224 and -1257 of the *Sox8* promoter while 2 sequences mapping to 2 distinct regions of *Mrhl* RNA, one mapping to the 5’ end of the lncRNA and the other to the 3’ end, were identified as TFO sequences. The 33nt region from the *Sox8* locus was split between 3 DNA oligos for downstream experiments while 2 RNA oligos were generated, each mapping to the two different predicted regions.

**Table 1.** Triplexator predictions: predictions with a score of 10 and above for regions within mrhl lncRNA (tfo start and end) and sox8 promoter (tts start and end)

| TFO start<br>(nucleotide<br>position within<br>mrhl) | TFO end<br>(nucleotide<br>position within<br>mrhl) | TTS start<br>(nucleotide<br>position from<br>TSS) | TTS end<br>(nucleotide<br>position from<br>TSS) | Score |
| --- | --- | --- | --- | --- |
| 391 | 404 | 1224 | 1237 | 11 |
| 389 | 400 | 1224 | 1235 | 10 |
| 389 | 404 | 1226 | 1241 | 12 |
| 389 | 404 | 1232 | 1247 | 12 |
| 389 | 401 | 1245 | 1257 | 10 |
| 2362 | 2374 | 1106 | 1118 | 10 |
| 2362 | 2374 | 1239 | 1251 | 10 |

EMSA was carried out to check for interaction between specific DNA and RNA oligonucleotides *in vitro*. A positive control TFO-TTS pair from within Meg3 lncRNA and its target TGFBR1 gene (Mondal et al, 2015) and a negative control TFO-TTS pair with no AG bias within the *Sox8* locus and *Mrhl* RNA were included for EMSA. Of the 5 TFO-TTS pairs probed, 2 combinations with *Mrhl* TFO1 as the RNA oligo, TTS2 and TTS3, showed a mobility shift corresponding to the formation of triplex. Triplex reactions were additionally subjected to RNase H digestion, which specifically degrades DNA:RNA duplexes to rule out any shift in mobility arising due to this hybrid duplex, in addition to RNase A digestion controls. While both TTS-TFO pairs showed a decrease in the intensity of signal with RNaseA treatment, only one *(Mrhl* TFO1-TTS2) was resistant to RNase H digestion (Fig 1B). Since the purine bases G and A participate in the formation of Hoogsteen base pairing, the G bases were mutated to C in TTS2 to look for the effect on triplex formation. We observed a loss in the mobility shift when the mutant oligos were used (Fig 1C). This pair was selected as a probable triplex-forming region between *Mrhl* and the *Sox8* locus.

To validate the results from EMSA, Circular Dichroism spectra for the TFO-TTS pairs TTS2-TFO1 and TTS2-NC TFO were recorded. The spectra recorded for the test oligos showed characteristics of a triplex with two minima- a strong negative peak at 210 nm and an additional negative peak at 240 nm (Fig 1E & 1F). Artificial spectra were generated by plotting the sum of the individual spectra for either TTS2 and TFO1 or TTS2 and NC TFO. When the artificial spectra were overlaid with the spectra obtained from the triplex reactions, an overlap in the spectra for the control oligo was observed but not for the specific TFO1 containing spectra indicating that the spectrum for the reaction containing control oligo is due to individual components (Fig 1E&F).

Further, the formation of triplex within the nucleus was investigated by using biotinylated TFO 1 or control TFO as bait to pulldown associated chromatin. Compared to the control oligo, TFO 1 pulldown fraction was significantly enriched for the *Sox8* TTS2 containing locus but not for a control region from the Beta-actin locus (Fig 1G). This enrichment was not significantly affected by RNaseH digestion thereby confirming that the TFO from within *Mrhl* lncRNA interacts with the *Sox8* locus through the formation of a DNA:DNA:RNA triplex.

### Triplex formation regulates Sox8 expression through PRC2 recruitment

Triplex formation by lncRNA can regulate genes present in its vicinity - either by activating or repressing gene expression. Specifically, triplex formation could act as a platform for the recruitment of DNMT3b which methylates the CpG island present in the vicinity, thereby leading to gene repression (Schmitz et al, 2010). Since the *Sox8* promoter and the TTS is located within a CpG island (supplementary Fig 2B), we wanted to investigate if lncRNA triplex formation leads to methylation of this CpG island to maintain *Sox8* in the repressed state. The methylation status was investigated in the murine cell line Gcl-spg, derived from the B-type spermatogonia, a cell line in which the role of *Mrhl* lncRNA has been extensively characterised, in the absence and presence of the Wnt ligand. Under control conditions, *Mrhl* is actively expressed and maintains *Sox8* in the transcriptionally repressed state and activation of the Wnt signalling pathway results in the downregulation of *Mrhl* lncRNA which activates *Sox8* expression (Akhade et al, 2014). To our surprise, no reduction in methylation levels was observed when *Sox8* transcription was activated. These results were corroborated by the results of methylation levels in mice testes of 7-day old and 21-day old mice (Supplementary Fig 2D). Testes of P7 mice have a predominant population of spermatogonial cells (cells without activation of Wnt signalling) while testes of P21 mice have a predominant population of spermatocytes (with activation of Wnt signalling). Much of the results pertaining to *Mrhl* lncRNA and *Sox8* expression and regulation in control and Wnt-signalling activated cells are also reflected in P7 and P21 mice testes respectively (Akhade et al, 2014, Akhade et al, 2016, Kataruka etal, 2017).

We have shown earlier that *Sox8* repression correlates with high levels of the repressive histone modification H3K27me3 in spermatogonia (Kataruka et al, 2017). So, we next checked for the presence of PRC2 at the promoter in the presence of *Mrhl* by performing ChIP for EZH2, a core subunit of PRC2 which enables PRC2- RNA interaction. The occupancy of Ezh2, which was observed under control conditions, was found to reduce upon activation of the Wnt signalling pathway in the Gcl-spg cells (Fig 2A).

**Fig 2:**
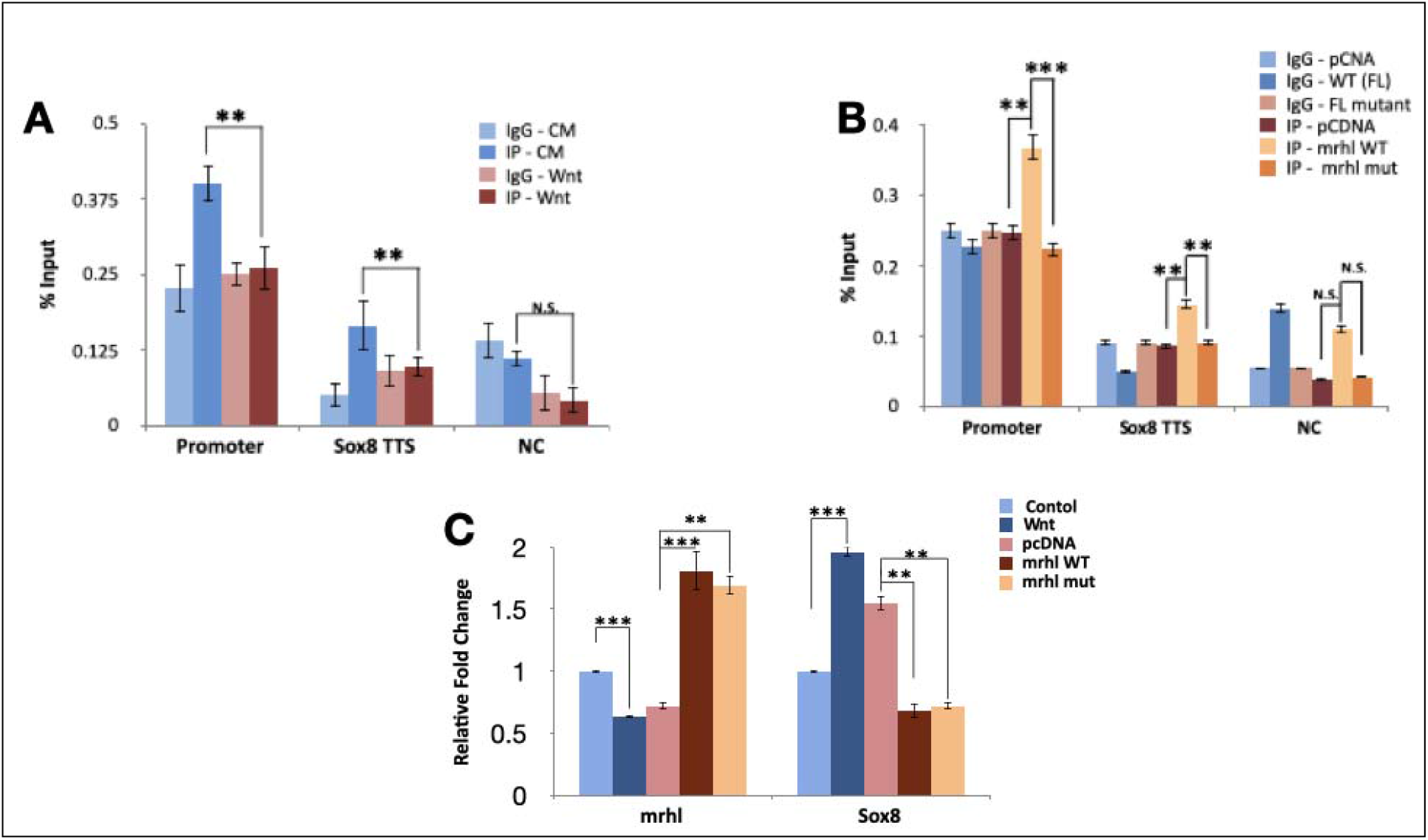
PRC2 occupancy at the Sox8 locus. **(A)** ChIP-qPCR for Ezh2 subunit of PRC2 at the Sox8 locus in Gcl-spg cells under control and Wnt signalling activated conditions. **(B)** ChIP-qPCR for Ezh2 performed, in cells transfected with either vector control, WT or TFO mutant constructs of Mrhl. **(C)** The expression levels of Mrhl and Sox8 in cells grown under control or Wnt activated conditions and cells transfected with vector only control Mrhl WT and Mrhl TFO mutant constructs with Wnt activation. Data in the graph has been plotted as mean ±S.D., N=3. *** P ≤ 0.0005, ** P ≤ 0.005, * P ≤ 0.05, N.S - Not significant (two-tailed Student’s t test)

Since multiple features including high GC-content of DNA, unique DNA conformations, presence of a guide lncRNA or the RNA:DNA:DNA triplex structure can recruit PRC2 to the target locus, we wanted to understand the contribution of the triplex formation by *Mrhl* lncRNA to the occupancy of PRC2 at the Sox8 locus. To this end, we attempted to rescue Ezh2 occupancy by expressing *Mrhl* lncRNA containing either the Wild-type (WT) TFO or mutated TFO. While WT *Mrhl* lncRNA expressed in *trans* in the presence of the wnt cue resulted in the rescue of Ezh2 occupancy, mutant TFO did not have a similar effect (Fig. 2B). Therefore, we conclude that the formation of triplex by *Mrhl* lncRNA at the *Sox8* locus recruits PRC2 to the *Sox8* locus. Furthermore, we also looked at the ability of TFO mutant *Mrhl* to rescue *Sox8* transcript levels. WT *Mrhl* was able to rescue to expression of *Sox8* to the levels comparable to control conditions (Fig 2C). Interestingly, the TFO mutant was able to transcriptionally repress *Sox8* similar to WT *Mrhl*. This indicates that triplex formation, while required for PRC2 recruitment, is not necessary to maintain *Sox8* in the repressed state (Fig. 2C) and points to additional role for *Mrhl* lncRNA in repressing *Sox8* transcription.

### CTCF and cohesin bind at the *Sox8* locus in the presence of lncRNA *Mrhl*

The p68-dependent gene repression of *Sox8* by *Mrhl* lncRNA is reminiscent of the role of lncRNA *SRA* and p68 at the imprinted *H19/Igf2* locus where the two molecules associate with architectural proteins CTCF and cohesin to silence *Igf2* gene on the maternal allele (Yao et al, 2010). We looked for potential CTCF and cohesin binding sites at the *Sox8* locus to check if a similar mechanism of gene repression was operating here. ENSEMBL database revealed that the transcription factor CTCF binds at a binding site present in exon 3 of the *Sox8* gene (Fig 3A). An inverse correlation between *Sox8* expression pattern and CTCF binding was observed across various tissues (supplementary fig 2B; activity at the *Sox8* gene locus is considered as an indication of *Sox8* expression) indicating that CTCF binding corresponds to the transcriptionally repressed state of the gene. The cohesin complex, too, binds at this region along with CTCF (supplementary 4A). CTCF binding is observed in those tissues in which *Mrhl* is expressed suggestive of cooperation between *Mrhl* and CTCF in the regulation of *Sox8* in multiple tissues (supplementary 4B).

**Fig 3:**
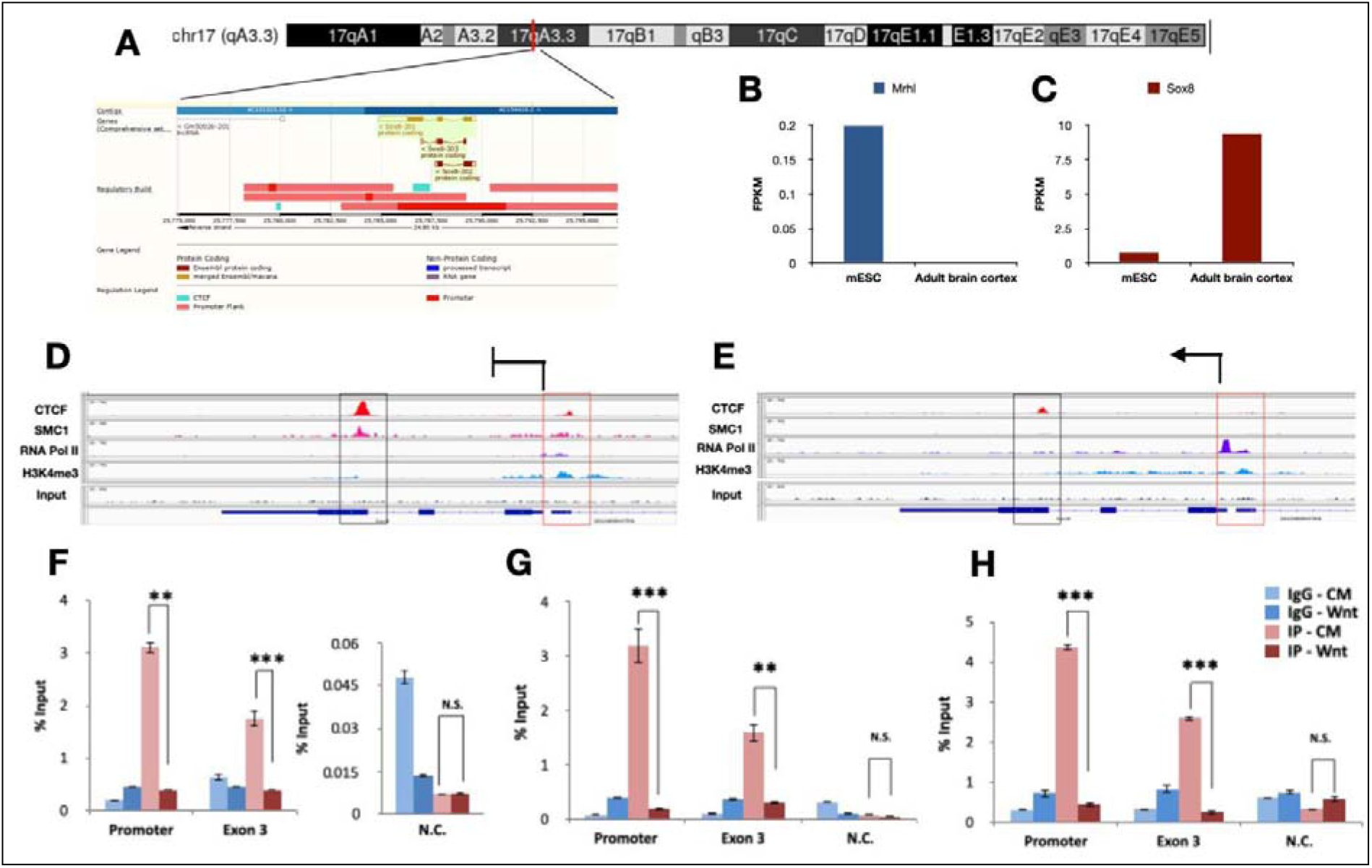
Occupancy of architectural proteins at the Sox8 locus in spermatogonia. **(A)** Sox8 locus harbouring CTCF binding site within exon 3. **(B)** Mrhl expression levels in mESC and adult brain cortex **(C)** Sox8 expression in mESC and adult brain cortex **(D)** Analysis of ChIP-seq datasets in mESC showing the presence of CTCF and cohesin (SMC1) at both exon 3 and the promoter **(E)** Analysis of ChIP-seq datasets in adult brain cortex showing reduced occupancy of CTCF and cohesin (SMC1) at exon 3 and promoter. Results for ChIP-qPCR for **(F)** CTCF, **(G)** Rad21 and **(H)** p68 showing their occupancy patterns at the promoter and exon3 of Sox8 gene in cells without and with Wnt activation. Data in the graph has been plotted as mean ±S.D., N=3. *** P ≤ 0.0005, ** P ≤ 0.005, * P ≤ 0.05, N.S - Not significant (two-tailed Student’s t test)

We then analysed publicly available ChIP-sequencing datasets to visualise the binding of the 2 architectural proteins at the *Sox8* locus in more detail. Due to the unavailability of required ChIP-seq datasets in spermatogonial cells, mouse embryonic stem cells and adult brain cortex were used as surrogate systems. *Mrhl* lncRNA is expressed in mESC while it isn’t expressed in the adult brain cortex (Pal et al, 2021). The inverse relationship between *Mrhl* and *Sox8* expression in these two tissues was confirmed by analysing publicly available RNA-seq datasets (Fig 3B&C). ChIP-Seq analysis for CTCF, SMC1 subunit of cohesin, RNA Pol II and H3K4mc3 to indicate transcriptional activity at the locus was carried out in these two surrogate systems. This confirmed the occupancy of CTCF within exon 3 of *Sox8* in mESC in which *Sox8* is not actively transcribed (low transcript levels in RNA seq and low relative levels of RNA polII at promoter and H3K4rne3 at the locus).

The same was found to be true for cohesin (the presence of SMC1 has been considered as evidence of cohesin occupancy). Interestingly, an additional peak for both CTCF and cohesin was observed at the promoter of *Sox8* very close to the *Mrhl* binding site (Fig 3D). Upon transcriptional activation (adult brain cortex) (higher transcript levels and relative occupancy of RNA polII at promoter and H3K4me3 at the locus), CTCF and cohesin occupancy were not observed at the promoter and the occupancy at exon3 was significantly reduced (Fig. 3E). This occupancy pattern was experimentally validated by performing ChIP for CTCF and Rad21 subunit of the cohesin complex in Gcl-spg cells without and with activation of the Wnt signalling pathway. A gene desert locus in chromosome 3 was chosen as a negative control for the ChIP-qPCR. To confirm that ChIP experiments were working, western blotting was carried out for the proteins (supplementary 4). The results from these experiments showed protein occupancy for both CTCF and Rad21 at both exon3 and *Sox8* promoter regions under control conditions. In the Wnt activated cells, CTCF and Rad21 binding at both positions in the *Sox8* locus was significantly reduced (Fig3F,3G). We knew from previous p68 ChIP experiments that it binds at the *Sox8* promoter at the *Mrhl* binding site. Our current p68 ChIP experiment showed significant enrichment of p68 at exon3 of *Sox8* in the Gcl-spg cells under Wnt inactivated conditions. This occupancy too reduced upon *Mrhl* downregulation by Wnt signalling activation (Fig. 3H).

To understand if the differential binding of these proteins at the *Sox8* locus was dependent upon *Mrhl* or was due to an indirect, downstream effect of the activation of the Wnt pathway, their occupancy was investigated in cells in which *Mrhl* was depleted by RNAi. ChIP was carried out in cells in which two different inducible lentiviral shRNA constructs targeting *Mrhl* or a non-target control construct were integrated (Fig. 4A). Similar to what was observed under Wnt induced conditions, occupancy of CTCF, Rad21 and p68, observed at both exon3 and promoter of *Sox8* in cells with non-target control, was depleted upon RNAi mediated *Mrhl* downregulation indicative of their *Mrhl* lncRNA dependent occupancy at the *Sox8* locus (Figs 4C, 4D & 4E).

**Fig 4:**
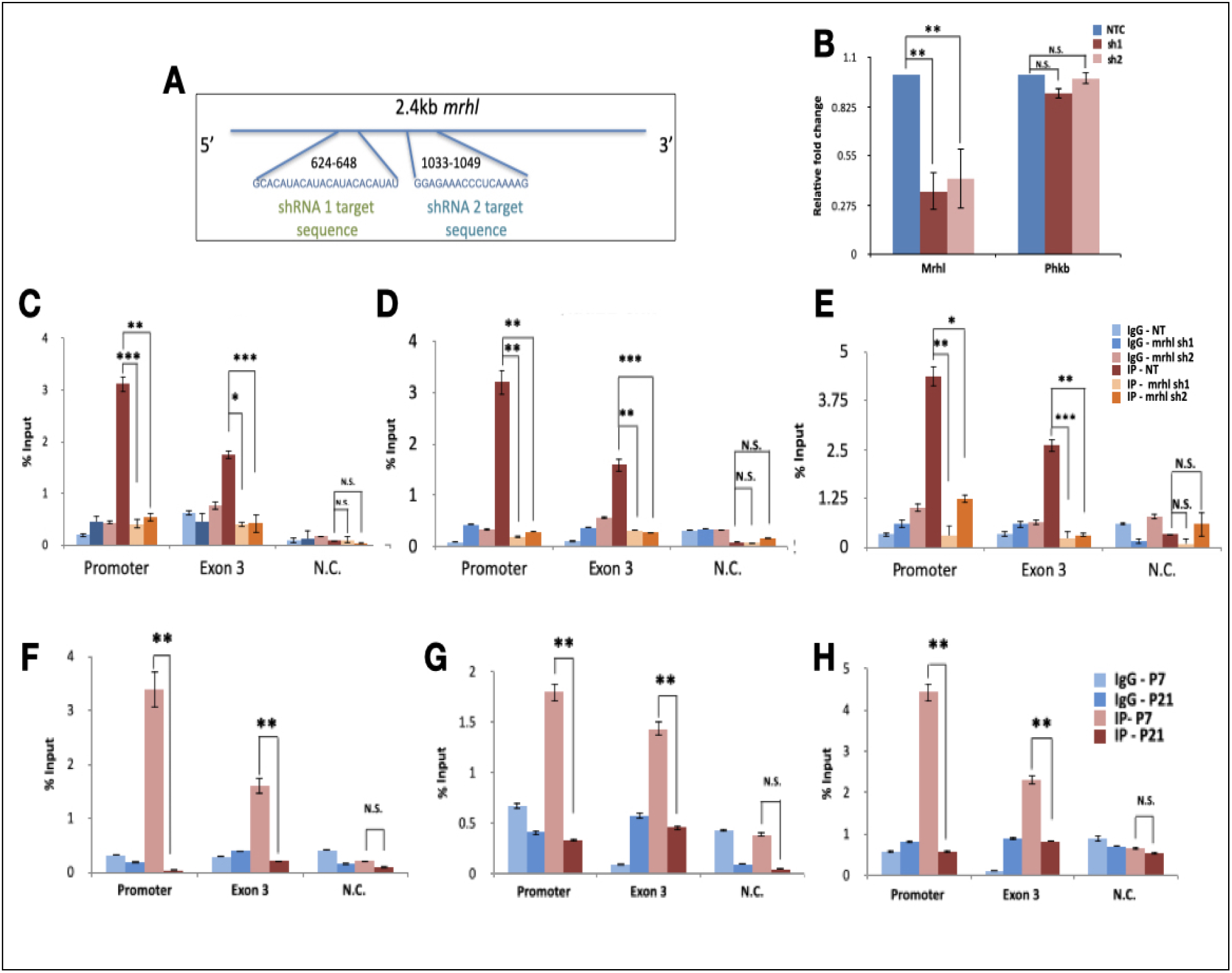
Architectural proteins at the Sox8 locus in Mrhl silenced cells and mice testes. **(A)** The two different regions within Mrhl targeted by the two shRNA **(B)** Mrhl silencing efficiency of the two shRNA-silencing efficiencies of ∼65% and ∼55% respectively were observed for Mrhl while the transcript levels of the host phkb gene were not perturbed significantly. Results for ChIP-qPCR for **(C)**CTCF, **(D)** Rad21 of cohesin and **(E)** p68 show significant reduction in occupancy of CTCF at both the promoter and exon 3 of Sox8 upon induction of silencing of Mrhl with both shRNA construct 1 and shRNA construct 2. Occupancy of **(F)** CTCF **(G)** Rad21 and **(H)** p68 is observed at the promoter and exon 3 of Sox8 locus in P7 mice testes and a significantly reduced occupancy is observed in P21 mice testes. Data in the graph has been plotted as mean ± S.D., N=3. *** P ≤ 0.0005, ** P ≤ 0.005, * P ≤ 0.05, N.S - Not significant (two-tailed Student’s t test)

We also investigated the status of occupancy of these proteins in the testes of mice in an attempt to address the biological relevance of the participation of CTCF and cohesin in *Sox8* gene regulation. Results of ChIP carried out for CTCF, Rad21 and p68 in P7 and P21 mouse testes corroborated with the results from control GCl-spg and Wnt induced/ RNAi induced *Mrhl* downregulated cells respectively (Figs 4F, 4G & 4H). The 3 proteins were found to be bound at the *Sox8* locus in the testes of 7-day old mice and a significant reduction in the occupancy was observed in 21-day old mice testes.

### YY1 binds at the *Sox8* locus upon *Mrhl* downregulation

CTCF and cohesin occupancy at the *Sox8* promoter was not seen in ENSEMBL database. *In silico* analysis using the Gene Promoter Miner tool was performed to look for probable interacting partners of CTCF that could be present at the *Sox8* promoter. The analysis of the promoter for transcription factor binding sites revealed the presence of a binding site for Ying-Yang 1 (YY1), a CTCF interacting transcription factor (Supplementary Fig 5A). Exon3 has characteristics of an enhancer-blocker or a silencer element, a type of cis-regulatory element of a gene which contributes to transcriptional silencing by contacting and recruiting repressive transcription factors such as CTCF to the promoter. If so, when not in the transcriptionally repressed state, the *Sox8* promoter is free to interact with an enhancer element. YY1 has been reported to bind at enhancers of genes and facilitate active transcription by enabling enhancer-promoter contact. As per the ENSEMBL database, two enhancer elements are present in the immediate vicinity of the *Sox8* gene, one upstream and one downstream of *Sox8*, and activity in both these elements correlated with transcriptional activity of *Sox8*. In an attempt to identify if YY1 enables the active transcription of *Sox8* by binding to its promoter and *Sox8* specific enhancer, we set out to identify potential enhancers for *Sox8* in the mouse spermatogonial cells. Multiple tissue-specific enhancers have been reported to regulate the expression of the members of the SoxE group of transcription factors. Specifically, the expression of *Sox9*, another essential transcription factor for sex determination and maintenance of mammalian testis, is regulated by multiple testis-specific enhancers, either synergistically or redundantly (Gonen et al, 2018). An early attempt to identify enhancers for *Sox8* identified 7 evolutionarily conserved regions in the vicinity of the gene with potential to act as enhancers. However, none of these elements acted as an enhancer in the embryonic gonad (Guth et al, 2010). Another study hinted at the presence of two regulatory elements downstream of the gene, specifically in cells of gonadal origin (Garcia-Moreno et al, 2019). Collating information from these different sources, we identified two regions downstream of *Sox8*, one 8kb downstream and another on 14kb downstream, as putative enhancers. Of these two regions, the element proximal to *Sox8* harbours one of the 7 evolutionarily conserved elements (E6).

We performed ChIP for the enhancer specific histone marks and for YY1 in Gcl-spg cells with and without activation of the Wnt signalling pathway to explore if (a) either one of the two putative elements gained enhancer specific histone modifications concomitant with *Sox8* transcriptional activation and (b) if YY1 could potentially be regulating *Sox8* expression by binding to the enhancer. Both the putative enhancer as well as the gene promoter regions had a significant increase in the levels of H3K4mel and H3K27ac marks upon Wnt induction (Figs 5A & B). Additionally, increased occupancy of YY1 at both of these enhancers and at the gene promoter was observed with Wnt activation (Fig 5C). The same trend was also observed in the RNAi mediated *Mrhl* knockdown cells. While the levels of H3K4mel and H3K27ac at both putative enhancer elements were low in cells with non-target control shRNA, the levels increase significantly in cells containing both *Mrhl* targeting shRNA (5D & E). Similarly, the occupancy of YY1 increased significantly at both the elements as well as the promoter upon *Mrhl* depletion (Fig. 5F) Finally, these observations were found to be biologically relevant since the testes of 21-day old mice showed significantly higher levels of H3K4mel and H3K27ac marks (Fig. 5G & 5H) as well as the occupancy of YY1 (Fig. 5I) at both the enhancer regions as well as the promoter of *Sox8* when compared to testes of 7-day old mice.

**Fig 5:**
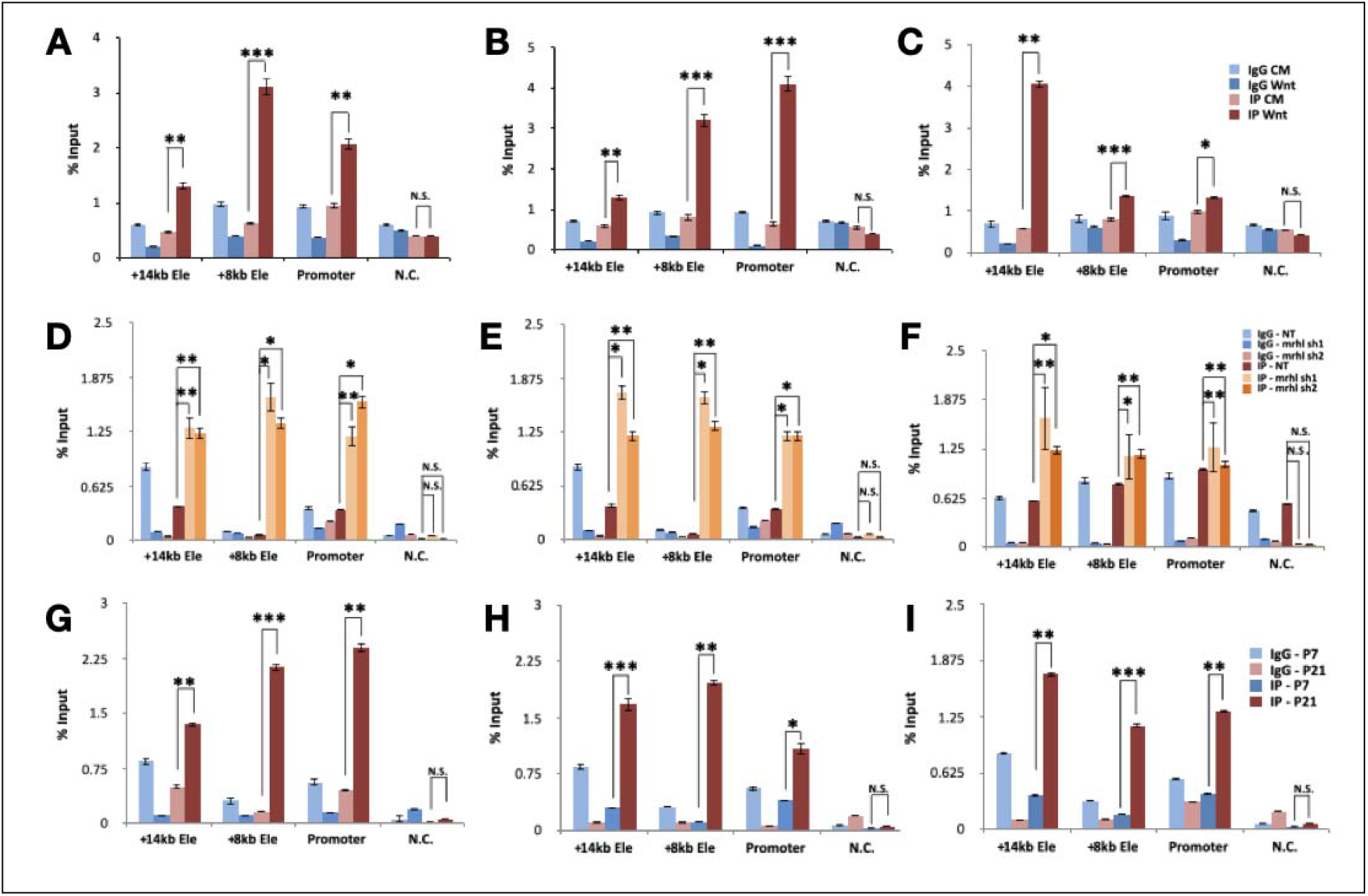
Enhancers at the Sox8 locus. Results from ChIP-qPCR experiment for **(A)** H3K27ac **(B)** H3K4mel and **(C)** YY1 show a significant increase in the levels of this modification with Wnt signalling induced Sox8 transcriptional activation at the promoter and both the enhancer elements. Results from ChIP-qPCR experiment for **(D)** H3K27ac, **(E)** H3K4mel and **(F)** YY1 show a significant increase in the levels of this modification with Sox8 transcriptional activation at the promoter and both the enhancer elements in Mrhl knockdown cells when compared to cells expressing non-target control shRNA. Results from ChIP-qPCR experiment for **(G)** H3K27ac, **(H)** H3K4mel and **(I)** YY1 show a significant increase in the levels of this modification in P21 mice testes when compared to P7 mice testes at the promoter and both the enhancer elements. Data in the graph has been plotted as mean ±S.D., N=3. *** P ≤ 0.0005, ** P ≤ 0.005, * P ≤ 0.05, N.S - Not significant (two-tailed Student’s t test)

### *Mrhl* mediates a looping switch at the *Sox8* locus

Many of the proteins bound at the promoter of *Sox8* including PRC2, p68, HDAC, Sin3a, and MAD-Max have been reported to interact with the proteins identified as bound within exon 3 of *Sox8* including CTCF and cohesin. We reasoned that occupancy of proteins such as p68 and Rad21 was detected at both the promoter and exon 3 because this protein complex was bringing the promoter and exon 3 in contact with each other through looping of chromatin at the *Sox8* locus. Silencer elements have been reported to repress gene transcription in precisely this manner - by binding to transcriptional repressors such as CTCF, contacting the gene promoter and preventing promoter-enhancer interaction (Ogbourne & Antalis, 1998). Additionally, we hypothesised that *Mrhl* down-regulation resulted in the dissociation of the promoter-silencer contact freeing the promoter to come in contact with the downstream enhancer element.

To validate the hypothesis, we looked at the chromatin looping status in the presence and absence of lncRNA *Mrhl* through Chromosome Conformation Capture (3C). From 3C performed in Gcl-spg cells with and without RNA knockdown, we observed that the *Sox8* promoter was indeed brought in contact with exon3 in a *Mrhl* dependent manner supporting our hypothesis. Interestingly, of the two enhancers identified downstream of *Sox8*, the interaction frequency between the promoter and the proximal enhancer (present 8kb downstream of TSS) was found to be higher in *Mrhl* knockdown cells while there was no significant difference in the interaction frequency between the promoter and more distal enhancer (14kb downstream of TSS) upon *Mrhl* depletion, suggesting an enhancer function for the proximal element in the mouse spermatogonial cellvs (Fig 6)

**Fig 6:**
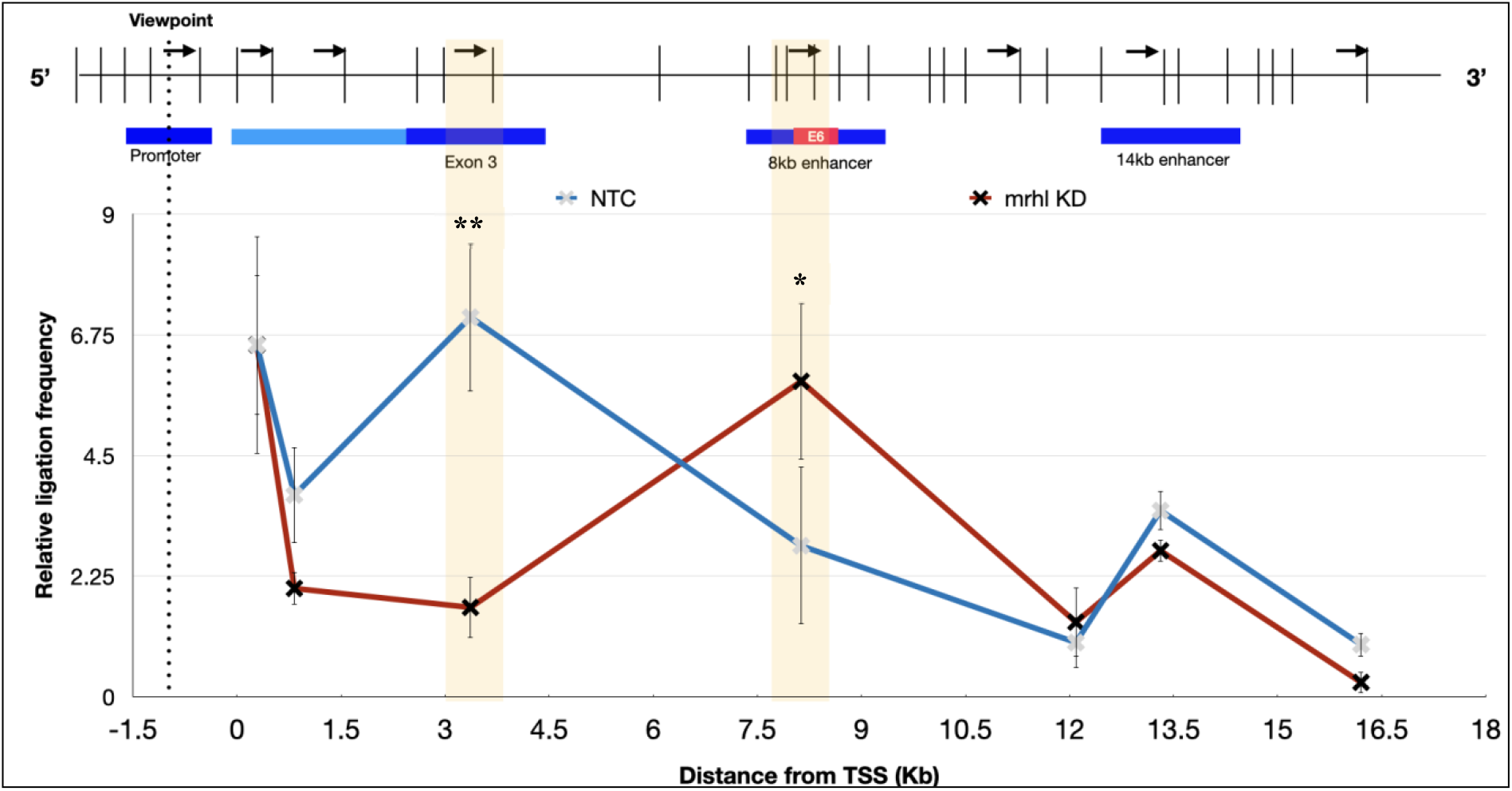
Chromatin looping at the Sox8 locus. 3C performed in Gcl-spg cells with non- target control shRNA (blue line) or Mrhl targeting shRNA (Red line). The schematic on top shows the positions of the DpnII sites (vertical bars) in the Sox8 locus relative to the genomic organisation. The black arrows indicate the 3C primer binding sites and their directionality. The viewpoint primer is within the promoter of Sox8 (indicated with dotted line). Graphs has been plotted as the relative ligation frequency (Y-axis) as a function of the distance from TSS in kb (X-axis). In the control cells, a peak in the relative ligation frequency is observed between promoter and exon 3 (highlighted with shaded box). In Mrhl knockdown cells, the peak at exon 3 is no longer observed but a peak at enhancer 8kb downstream of TSS (highlighted with shaded box) is observed indicative of interaction of the promoter with this segments of DNA. Data in the graph has been plotted as mean ± S.D., N=3. *** P ≤ 0.0005, ** P ≤ 0.005, * P ≤ 0.05, N.S - Not significant (two-tailed Student’s t test)

## Discussion

A member of the SoxE group, *Sox8* is essential for the maintenance of male fertility as *Sox8* null mice show progressive degeneration of spermatogenesis resulting in infertility (O’Bryan et al, 2008). Specifically, *Sox8* expression in Sertoli cells is essential for germ cell differentiation (Singh et al, 2009). Most of the previous studies have focussed on understanding the role of *Sox8* in Sertoli cells in mammalian testes. Studies from our group were the first to not only report the expression of *Sox8* in spermatogenic cells but also explore the potential role of this transcription factor in meiotic commitment, likely via the master regulator of meiosis in spermatogenesis, *Stra8* (Kojima et al, 2019). In this context, it was of importance to study the detailed molecular events during the regulation of *Sox8* by *Mrhl* lncRNA.

The formation of a DNA:DNA:RNA triplex is a mechanism of interaction that is common to many chromatin bound lncRNAs such as *Meg3, PARTICLE, HOTAIR* and *KHPS1* (Mondal et al, 2015, O’Leary et al, 2015, Kalwa et al, 2016, Blank-Giwojna et al, 2019). In the current study, we show that *Mrhl* too interacts at the *Sox8* locus directly with the chromatin through the formation of DNA:DNA:RNA triplex. *Mrhl* lncRNA harbours multiple potential triplex forming regions within it. The *in silico* predictions for the *Sox8* locus suggested two different regions to have triplex forming potential - one mapping to the 5’ end and another towards the 3’ end. While the sequence towards the 5’ end participates in triplex formation at the *Sox8* locus in the Gcl-spg cells, it is possible that the sequence towards the 3’ end forms triplex in a different context. The predictions from Triplexator using different genomic regions such as Pou3f2, Runx2 or FoxP2 (Pal et al, 2021), show that other regions within *Mrhl* lncRNA, too, have the potential to form triplex. It is likely that *Mrhl* interacts at multiple other loci through the formation of DNA:DNA:RNA triplex through different TFOs present within it in a context dependent manner.

### PRC2 in Sox8 gene regulation

Many genes are repressed through methylation of CpG rich DNA at their promoters. Further, triplex formation by the ncRNA pRNA acts as a platform for the recruitment of the DNA methyltransferase DNMT3b at the rDNA promoter which goes on to methylate the DNA at the gene promoter, thereby repressing transcription (Schmitz et al, 2010). The presence of a 1.3kb long CpG island encompassing the promoter of *Sox8* suggested s probable mechanism of gene repression through the methylation of this CpG island. However, no reduction in methylation levels was observed experimentally corresponding to *Sox8* activation in either the mouse spermatogonial cell line Gcl-spg upon Wnt induction or in 21-day old mouse testes suggesting that DNA methylation is not the mechanism of epigenetic repression of *Sox8*.

The presence of high levels of H3K27me3 repressive histone mark in the *Sox8* transcriptional repressed state (Kataruka et al, 2017) was indicative of the presence of PRC2 at the *Sox8* locus. PRC2 is the multi-protein complex responsible for catalysing the methylation of H3K27. A common feature of the mammalian PRC2-binding region is the presence of CpG islands (CGIs) and more specifically, CpG-rich regions which are adjacent to the TSS of silenced genes. Multiple studies indicate that high-density DNA methylation seems to be mutually exclusive with PRC2 since most of the CGIs or CG-rich regions occupied by PRC2 are hypomethylated (Yang and Li, 2020). In agreement with these studies, the levels of methylation are lower at the promoter of Sox8, which is situated within the CpG island, in the *Sox8* transcriptionally repressed state than in the active state.

Another factor influencing PRC2 binding to target loci is its interaction with RNA molecules. It is now believed that lncRNA interaction, including those with PARTICLE and Meg3, could be one of the mechanisms by which PRC2 gains target specificity (Mondal et al, 2015, O’Leary et al, 2016). Different subunits of PRC2 recognise and bind to different secondary structures/ DNA sequences through which they get targeted specifically to genomic loci. For instance, unmethylated GCG trinucleotide motif showing an unwound DNA helix can specifically recruit PRC2-MTF2 while the Suz12 subunit has been reported to bind to the two-hairpin motif present in RNA molecules. PRC2 subunit JARID2 preferentially binds to GC rich DNA sequences (Yang and Li, 2020). At the *Sox8* locus, multiple possible modes of recruitment of PRC2 exist, namely, the presence of a hypomethylated CpG island, the presence of *Mrhl* lncRNA and also the formation of triplex by *Mrhl* lncRNA. Through functional rescue with wild type and TFO mutant *Mrhl*, we have shown that triplex formation indeed recruits PRC2 to the *Sox8* locus.

### CTCF and cohesin in Sox8 gene regulation

CTCF in mammals is the master architectural protein and along with cohesin, has been implicated in organizing chromatin architecture at different genomic scales from chromatin loops at the scale of a single locus to the organization of TADs. Using a combinatorial approach, we show the *Mrhl* dependent occupancy of CTCF and cohesin at the *Sox8* locus along with the DEAD-box RNA helicase p68. The popular loop-extrusion model of chromatin loop formation proposes that the cohesin protein complex slides along chromatin forming a growing loop until it meets two CTCF molecules bound with convergent orientation. This prevents cohesin from sliding further. Preliminary *in silico* analysis suggests the presence of CTCF binding sites both at the promoter and within exon3 of *Sox8* (supplementary table 1). A limitation of the prediction software utilised for this study is that it does not exhaustively predict the presence of all CTCF binding sites present within the sequence but only the sequence with the highest score. Heterogeneity is observed in CTCF binding motifs. In each species, the CTCF binding profile is composed of substantial numbers of both deeply conserved and evolutionarily recent sites. CTCF binding sites at TAD boundaries are highly conserved across species while evolutionarily recent sites play a role in modulating gene regulation (Kentepozidou et al, 2020). Further, cell-type specific CTCF bound sites have also been reported to have a varied binding motif as compared to constitutively bound sites (Essien et al, 2009). This is a likely explanation for why CTCF binding site was not predicted to be present by the GPMiner tool (Fig. 5A).

CTCFL is a testis-specific paralog that is expressed only transiently in pre-meiotic male germ cells together with CTCF and the two paralogs compete for binding at a subset of the CTCF binding sites (Nishana et al, 2020). CTCFL functions as a transcription factor and does not have a role in chromatin organisation since it can’t anchor cohesin to chromatin like CTCF can (Pugacheva et al, 2020). ChIP qPCR using CTCF specific antibody in the spermatogonial cells performed by us (supplementary figure 7) further confirms that CTCF and not CTCFL is bound at the *Sox8* locus.

DNA methylation at the DMR regulating the imprinting at the H19/Igf2 locus prevents the binding of CTCF to its cognate binding site within the DMR. The slight increase in the methylation at the CpG island of the *Sox8* promoter (supplementary 1D) upon its transcriptional activation may possibly serve the same purpose.

### Silencer and enhancer elements in Sox8 gene regulation

In addition to promoters, silencers/insulators and enhancers together make up *cis*-regulatory elements (CREs) of a gene. H3K27me3 mark enrichment has been found to be enriched within silencer elements. Most H3K27me3^+^ silencer elements are also DNase I Hypersensitive and have binding sites for ubiquitous repressors such as CTCF, SMAD group of proteins and tissue specific TFBS (Huang, D. et al, 2019). Additionally, many H3K27me3-DHS coincided with active histone modifications such as H3K4mel and H3K27ac. The element within exon 3 has many of these characteristics. In addition to the occupancy of CTCF and cohesin within this genomic region, ENSEMBL database suggests that this element is DNase hypersensitive and shows the presence of both H3K4mel and K3K27me3. The results of the 3C experiment further indicate that this element contacts the gene promoter in the transcriptionally repressed state. Taken together, this evidence supports the ‘silencer’ function of exon 3 of *Sox8*.

We have identified two putative enhancers for *Sox8* in spermatogonial cells located downstream of the gene. We observe activity at both these enhancer elements upon *Mrhl* knockdown as evidenced from enhancer specific histone modification ChIP experiments. Only one of these two regions, the enhancer present 8kb downstream, contacts the promoter of *Sox8* as observed from the 3C experiment and is likely to drive the expression of *Sox8* in meiotically committed spermatogonia. This enhancer harbours within it one of the evolutionarily conserved elements, E6 (Guth et al, 2010), as a putative enhancer. However, this does not mean that the other enhancer element has no role to play in regulating *Sox8* expression or that the E6 harbouring enhancer is the sole enhancer regulating the expression of *Sox8*. Further, this study has been performed in embryonic gonads and gives us no information on the postnatal activity of this regulatory element. The possibility that E6 harbouring enhancer may not be a testis-specific enhancer exists.

*Sox9* is regulated by multiple tissue specific enhancers and in the testis alone, is regulated by multiple enhancers acting either redundantly or synergistically (Gonen et al, 2018, Croft et al, 2018). Our study has focussed on characterising the regulatory elements of *Sox8* in a genomic region of 25-30kb only. Extensive characterisation including genomic deletion of the regulatory elements in a larger region is required to identify and better understand the possible interplay between various enhancers in regulating *Sox8* expression. Such characterisation is beyond the scope of the current study. Further, the possibility of *trans-*interactions regulating Sox8 expression has not been explored.

### YY1 in Sox8 gene regulation

Of all the regulatory proteins identified at the *Sox8* locus, YY1 is the only one with contradictory functional roles. YY1 can act both as a transcriptionally activator or transcriptional repressor in a context dependent manner. As an architectural protein too, YY1 can mediate the formation of chromatin loops which can either have gene repressive or activating outcomes.YY1 dimerises with CTCF to mediate chromatin loop formation to repress E6 and E7 oncogenes of the human papillomavirus genome in infected cells (Pentland et al, 2018). At the same time, YY1 binds to promoter-proximal elements and active enhancers and forms dimers that facilitate the interaction of these DNA elements (Weintraub et al, 2017). Thus YY1 at the *Sox8* locus was a wildcard that could be involved in either function. However, the results from the ChIP experiments clearly indicated the association of YY1 at the regulatory elements only in the *Sox8* transcriptionally active state. Further, the occupancy of YY1 at the promoter and enhancer elements suggested a role for it in facilitating the interaction of the enhancer with the promoter and this has been validated by chromosome conformation capture.

*Mrhl* lncRNA possible possesses multiple functional domains within it. At the *Sox8* locus, a region from the 5’ end of the lncRNA participates in triplex formation. Results from previous work suggest that a region towards the 3’ end of *Mrhl* is involved in its interaction with p68 (unpublished data). The gene regulatory function is an outcome of the combinatorial function of all the different domains of *Mrhl*. Further, many lncRNAs involved in 3D genome organisation have been reported to interact directly with CTCF. It remains to be seen whether *Mrhl* lncRNA also physically interacts with CTCF directly. *Mrhl*, then, can be categorised as scaffold lncRNA which functions to bring together multiple regulatory proteins at the target locus.

In the current study, we have demonstrated that the lncRNA *Mrhl* is involved in mediating chromatin looping at the *Sox8* locus in association with the architectural proteins CTCF and cohesin to maintain the gene in the transcriptionally repressed state. The downregulation of *Mrhl* lncRNA results in a rearrangement of the looping interaction at the locus whereby the promoter-silencer contact gives way to a promoter-enhancer contact mediated by YY1. Further, *Mrhl* forms a DNA:DNA:RNA triplex at the distal promoter of *Sox8* that is required for the recruitment of PRC2 which then tri-methylates H3K27 at *Sox8* gene locus (summarised in fig. 7).

**Fig 7:**
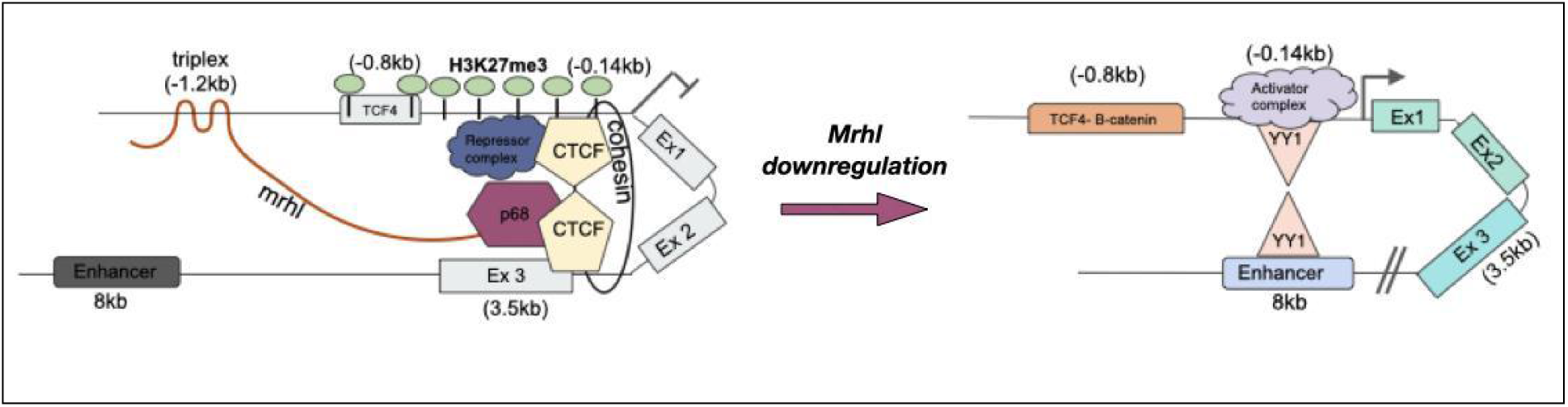
Model summarising the regulation of *Sox8* in the B-type spermatogonial cells.

The mechanism of silencing at the *Sox8* locus by *Mrhl* lncRNA via triplex formation, PRC2 recruitment, and the involvement of CTCF, cohesin and p68 fits into the growing theme of gene silencing mechanism by lncRNAs. Stating that *Mrhl* associates with this protein complex to mediate the formation of a repressive loop is a simplistic view of events. Taking into account the very large size of the repressive complex made up of CTCF, cohesin complex, Sin3a, HDAC1, Mad-Max transcription factor dimer, p68, PRC2 and *Mrhl*, it would be more realistic to state that *Mrhl* creates a repressive environment around the *Sox8* locus.

In the recent update of ENSEMBL, the human *Sox8* locus can be seen to not only contain a conserved CTCF binding site in exon 3 but also has binding of many of the regulatory protein observed at the mouse *Sox8* locus. While the data is indicative of a conserved regulatory mechanism, the involvement of a lncRNA in the regulation of *Sox8* in humans remains to be seen.

In summary, we have delineated the detailed molecular mechanism of regulation of Sox8 gene expression by Mrhl lncRNA which has significant implications towards our understanding the role of Sox8 in male germ cell differentiation, particularly meiotic commitment of B-type spermatogonia.

## Materials and Methods

### Cell lines and reagents

Gcl-spg cell line (CRL-2053) was obtained from ATCC. L-control cell line (ATCC CRL-2648) and L-Wnt3A cell line (ATCC CRL-2647) were kind gifts from Dr. Jomon Joseph (NCCS, India).

All chemical reagents were purchased from Sigma-Aldrich and were of analytical reagent (AR) grade. Lipofectamine 2000 (11668027) and Protein A dynabeads (10002D) were purchased from In vitro gen. cDNA synthesis reagents were procured from Thermo Fisher Scientific. DpnII restriction enzyme (R0543), DNase I (MO303) and T4 DNA ligase (M0202S) were purchased from New England BioLabs. Mrhl shRNAs (custom synthesised) and non-target control (SHC332) were procured from Sigma-Aldrich. qPCR was performed using BioRad’s CFX96 machine. DNeasy Blood and Tissue kit (69504), EpiTect II DNA Methylation Enzyme Kit (335452) and EpiTect methyl II PCR system (EPMM104707-1A) for CpG methylation analysis was procured from Qiagen. [γ-32P]ATP was sourced from

List of antibodies that were used are as follows with the manufacturer and catalog number in parenthesis are: CTCF (AbCam; ab 188408), (Cell Signaling Technology; 3418), Rad21 (AbCam; ad9920), p68/DDX5 (Cusabio; CSB-PA003685), YY1 (Diagenode; C15410345), H3K4mel (Diagenode; C15410037), H3K27ac (Diagenode; C15410174), Ezh2 (Diagenode; C15410039).

### Cell Culture and preparation of Control and Wnt3A conditioned media

Gc1-spg cells were cultured in Dulbecco’s modified Eagle’s Media (DMEM) supplemented with 10% Fetal Bovine Serum (FBS) and Penicillin/Streptomycin.

Preparation of control and Wnt3A conditioned media was done as per manufacturer’s (ATCC) instructions. The collected media was centrifuged at 500Xg for 5 min and used after filter sterilisation.

Gc1-spg cells were grown in control or Wnt conditioned media for 72 hours for Wnt induction. All cell-lines were checked for Mycoplasma contamination every 2 months

### Generation of stable knockdown lines

All shRNA plasmids were transfected at a concentration of 1.5 μg/ml Lipofectamine 2000 at 70% cell confluence. To select for positive transfectants, cells were grown in selection media containing puromycin at a final concentration of 3ug/ml for 72 hours. To induce shRNA expression, cells were grown in complete media in the presence of 0.5mM IPTG and 2.5ug/ml puromycin for 96 hours.

### Cloning

Full length WT mrhl and TFO mutant mrhl was cloned into pCDNA3.1 vector between the HindIII and BamHI sites. Clones were confirmed by Sanger sequencing.

### Preparation of the mice testicular samples

The testicular samples prepared were harvested from BALB/C mice in the ages groups of 7dpp and 21dpp. The dissected testis was decapsulated in PBS (pH 7.4) on ice by removing the tunica albuginea. The seminiferous tubules were released into PBS (pH7.4) and subjected to homogenization to procure a single-cell suspension.

### Chromatin immunoprecipitation

ChIP was performed as previously described (Kataruka et al, 2017). Briefly, crosslinked cells were resuspended in SDS lysis buffer (1% SDS, 10 mM EDTA, 50 mM Tris-HCl). This was followed by sonication of the lysate to enrich for chromatin in the size range of 200-600bp.After removal of debris by contrifugation, the lysate was incubated with either 3-5ug specific antibody or corresponding amount of isotype control for immunoprecipitation overnight. The immune complexes were allowed to bind to protein A Dynabeads and the beads bound by immune complexes were subjected to washes with low-salt buffer (0.1% SDS, 1% Triton X-100, 2 mM EDTA, 20 mM Tris-HCl, 150 mM NaCl), high-salt buffer (0.1% SDS, 1% Triton X-100, 2 mM EDTA, 20 mM Tris-HCl, 500 mM NaCl), LiCl wash buffer (1%NP-4O, 1% Sodium deoxycholate, 1mM EDTA, 10mM Tris-Cl pH 8.0) and T.E (10mM Tris-Cl pH 8.0, 1mM EDTA). The beads were then either processed directly for western blotting or the immunoprecipitated material was eluted from the beads by adding elution buffer (0.1 M NaHCO3, 1% SDS). The supernatant treated with Proteinase K (Life Technologies) and crosslinks were reversed. Eluted DNA was used for real-time PCR.

### Western blotting

After SDS-PAGE and transfer, the membranes were blocked in 5% skimmed milk for 1 hour at room temperature and then incubated with the respective primary antibody dissolved in 1% skimmed milk prepared in 1XPBS with 0.1% Tween 20 (0.1% PBST). The membrane was then washed with 0.1% PBST and incubated with the respective secondary antibody dissolved in 1% skimmed milk for an hour at room temperature. Washes were then given using 0.1% PBST, and the blot was developed using Millipore’s Immobilion Forte Western HRP substrate and the image was captured on BioRad’s ChemiDoc.

### Circular Dichroism spectroscopy

CD-spectra were recorded on a Jasco 500A spectropolarimeter. Each spectrum is the average of 4 consecutive spectra (independent replicates) and baseline-corrected with a spectrum of pure buffer. CD-spectra were recorded on *mrhl* ssRNA TFO, NC ssRNA TFO and the dsDNA oligos separately as well as on a 1:1 mix of the two in 1X triplex forming buffer (10mM Tris pH 7.5, 25mM NaCl and 10mM MgC12). The mixed samples or individual RNA and dsDNAs were heated to 70°C for 10 minutes and slowly cooled to room temperature and incubated at room temperature for 2 hours. The samples were incubated overnight at 4°C. The measurements were performed at 5 °C The data presented in the spectra is the Molar Ellipticity given based on the concentration of nucleotides in the samples.

### Triplex Electrophoretic Mobility Shift Assay

Electrophoretic mobility shift assay was performed as per protocol of Mondal et al (Mondal et al, 2015). Double stranded Oligonucleotides were end-labeled with T4 polynucleotide kinase in the presence of [γ-32P]ATP and purified using G-25 columns (GE Healthcare). To remove secondary structures present in them, RNA oligonucleotides were heated at 70°C and incubated on ice. Binding reaction was carried out in using labeled dsDNA oligonucleotides (corresponding to 10,000cpm), 1X Triplex forming buffer and 50 times molar excess of RNA oligonucleotides and incubated for 6 hours at room temperature. In control assay, triplex reaction was treated with either 5 units of RNase H (NEB) or 10εg of RNase A (Life Technologies) for 20min. Triplex formation was monitored on 20 % polyacrylamide TBE gel in 1X TBE buffer supplemented with 8 mM MgC12. The gel was dried and exposed to X-ray films.

### *In vitro* triplex pulldown assay

Nuclei from Gc1-spg cells were prepared by resuspending the cells in 1X nuclei isolation buffer (40mM Tris-HCl pH7.5, 20mM MgCl2, 4% tritonX-100, 1.28M sucrose). 10 μM psoralen-biotinylated TFO1 or NC TFO RNA oligonucleotides (purchased from Sigma-Aldrich) were incubated with the Gc1-spg nuclei for 2.5 hours at 30°C in 1X Triplex forming buffer followed by 10 minutes of UV treatment to induce photo-adduct formation. Nuclei were lysed using Bioruptor (10 cycles, 30 sec ON and 30sec OFF, Diagenode). The Supernatants were incubated with 50μl streptavidin-magnetic beads at 30°C. In case of RNase H control reaction, the supernatants were treated with 15 units of RNase H for 20 min before streptavidin-magnetic beads were added. Following beads capture, the beads were washed in 1X Triplex forming buffer to remove the non-specifically bound DNA fragments and then beads were resuspended in DNA isolation buffer. Resuspended beads bound to DNA-RNA triplex were treated with RNase A (20 μg) for 30 minutes followed by Proteinase K treatment. Precipitated DNA was used as template for qPCR.

### Chromosome conformation capture assay

BAC plasmids containing the Sox8 locus (RP23-70O24) and the control Ercc3 locus (RP23-148C24) (BACPAC resources, California) were purified by Alkaline Lysis method. The plasmids were mixed in equimolar ratio and 20μg of mixed plasmid was subjected to restriction digestion with the enzyme Sau3AI (NEB, Catalog number: R0169) (Isoschizomer of DpnII) overnight at 37°C. DNA was precipitated and resuspended in ligation master mix (1X NEB Ligation buffer, 0.8% Triton X-100, 0.5X BSA, 1600U of T4 DNA ligase (NEB, Catalog number: M0202). The reaction was incubated at 21 °C for 4 hours. Ligated DNA was precipitated and used as a template for PCR.

Contact library was generated as described by Mumbach et al (Mumbach et al, 2016) with modifications. Briefly, nuclei were isolated from crosslinked cells by resuspending cells in cell lysis buffer (10mM Tris-Cl pH 8.0, 1.5mM MgCl2, 10mM KCl,0.5mM DTT, 0.05% NP-40, IX mPIC and 0.2μM PMSF) and incubating on ice for 30 minutes. Pelleted nuclei were washed once with cell lysis buffer and permeabilised using 0.7% SDS by incubating at 62°C for 15 minutes. Triton X-100 was added to a final concentration of 10%. 50μl of 10X DpnII buffer and 375U of DpnII (NEB, Catalog number: R0543) restriction enzyme was added and the reaction was incubated overnight at 37°C with shaking at 900rpm. The reaction was heat inactivated at 62°C for 20 minutes. In-situ ligation was carried out by adding ligation master mix and incubating at 21 °C for 4 hours with shaking. Nuclei were pelleted and resuspended in SDS Lysis Buffer. The lysate was subjected to proteinase K treatment and reverse-crosslinking overnight at 65°C. DNA precipitated was then used as template for PCR. The relative frequency of interaction was calculated as described by Naumova et. al, (Naumova, N. et al, 2012) from agarose gel images.

### CpG methylation assay

The methylation status at the Sox8 promoter was scored for using the EpiTect methyl II PCR kit from QiaGen according to manufacturer’s instructions.

### Triplexator prediction

To identify all putative triplexes that can form between *mrhl* lncRNA and Sox8 promoter, analysis was run with default parameters (Buske et al, 2012) to identify TFO-TTS pairs in single-strand and duplex sequences.

### ChIP-sequencing data analysis

Raw FASTQ files were downloaded from the NCBI GEO repository, and were re-analysed to generate the aligned files for the visualisation of regions of interest. All aligned files were aligned to mouse genome (mm10) using Bowtie2 (Langmead and Salzberg, 2012) and then sorted, indexed, made free from PCR duplicates using Samtools (Li et al, 2009). Aligned files were loaded in the IGV genome browser to visualise the enrichment of peaks at the regions of interest. Peak calling was done with the MACS2 pipeline (Feng et al, 2012).

### RNA-sequencing data analysis

Raw FASTQ files were downloaded from the NCBI GEO repository, and were re-analysed with the TopHat (Kim et al, 2013) and Cufflinks (Trapnell, C. et al, 2010) pipeline. Aligned files were loaded on the IGV genome browser (Robinson et al, 2011) to visualise the gene expression enrichment.

### IACUC approval

Experiments were performed using mice testes. The institution (JNCASR) has obtained IACUC approval for research involving use of mice.

---

**Supplementary Table 1:**
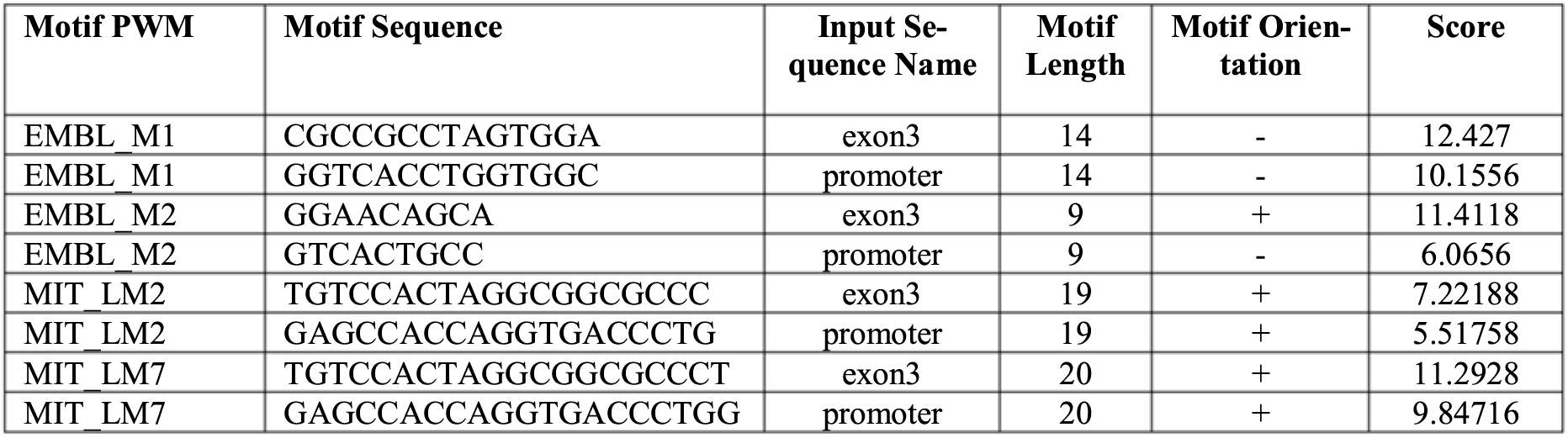
Predicted CTCF binding site from CTCFBSDB 2.0 database based in the DNA sequences of Sox8 promoter and exon 3. All results with a score higher than 3 have been listed in the table

**Supplementary Table 2:** List of datasets analysed for the present study

| ChIP-seq datasets | GEO Accession number |
| --- | --- |
| mESC CTCF | GSM723015 |
| mESC SMC1 | GSM560341 |
| mESC RNA Pol II | GSM723019 |
| mESC H3K4me3 | GSM723017 |
| mESC Input | GSM723020 |
| Mouse brain cortex CTCF | GSM722631 |
| Mouse brain cortex SMC1 | GSM1838869 |
| Mouse brain cortex RNA PolII | GSM722634 |
| Mouse brain cortex H3K4me3 | GSM722633 |
| Mouse brain cortex input | GSM722635 |
| RNA-seq datasets |  |
| mESC | GSM723776 |
| Adult Brain cortex | GSE96684 |

**Supplementary Table 3:**
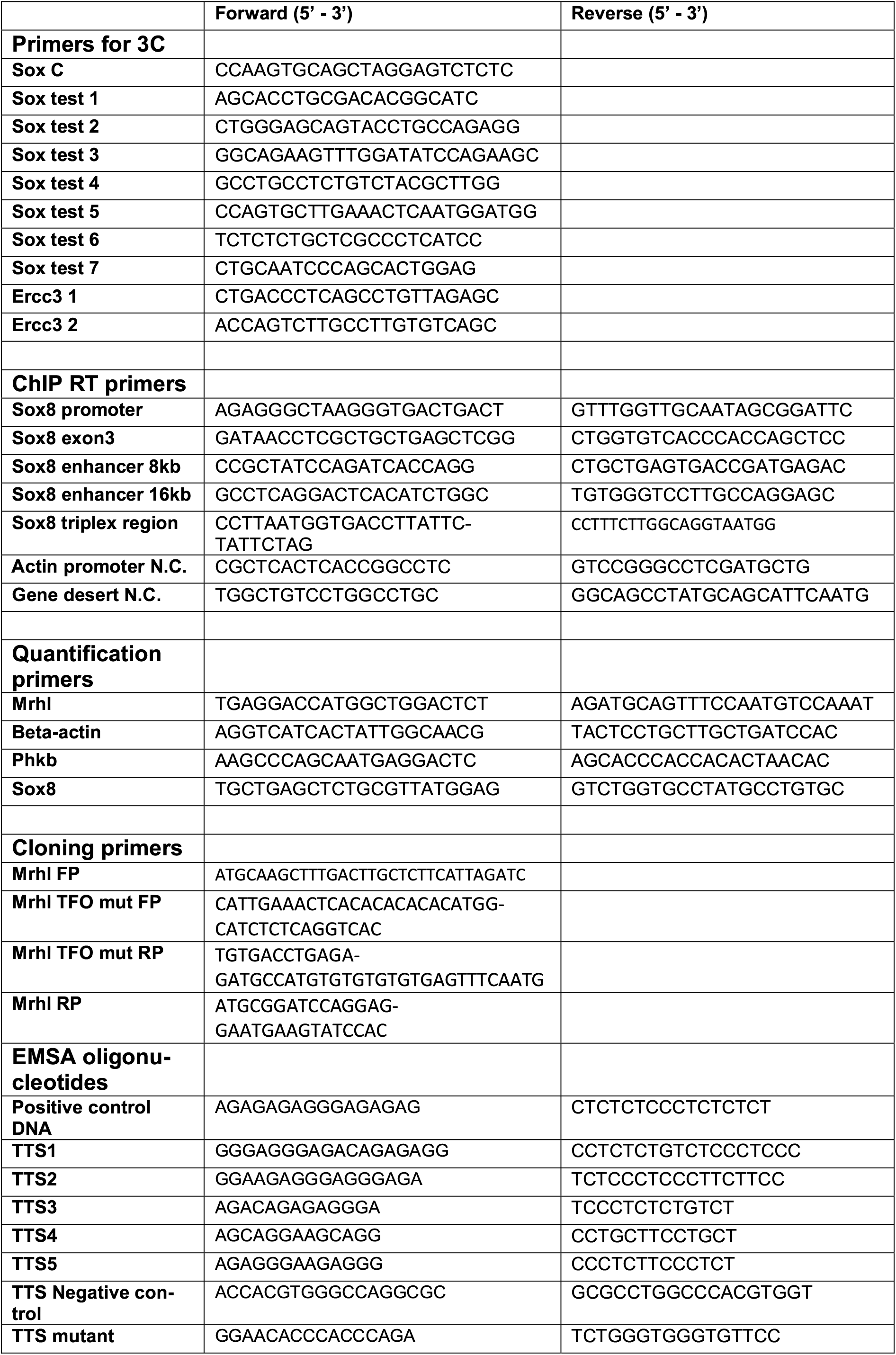
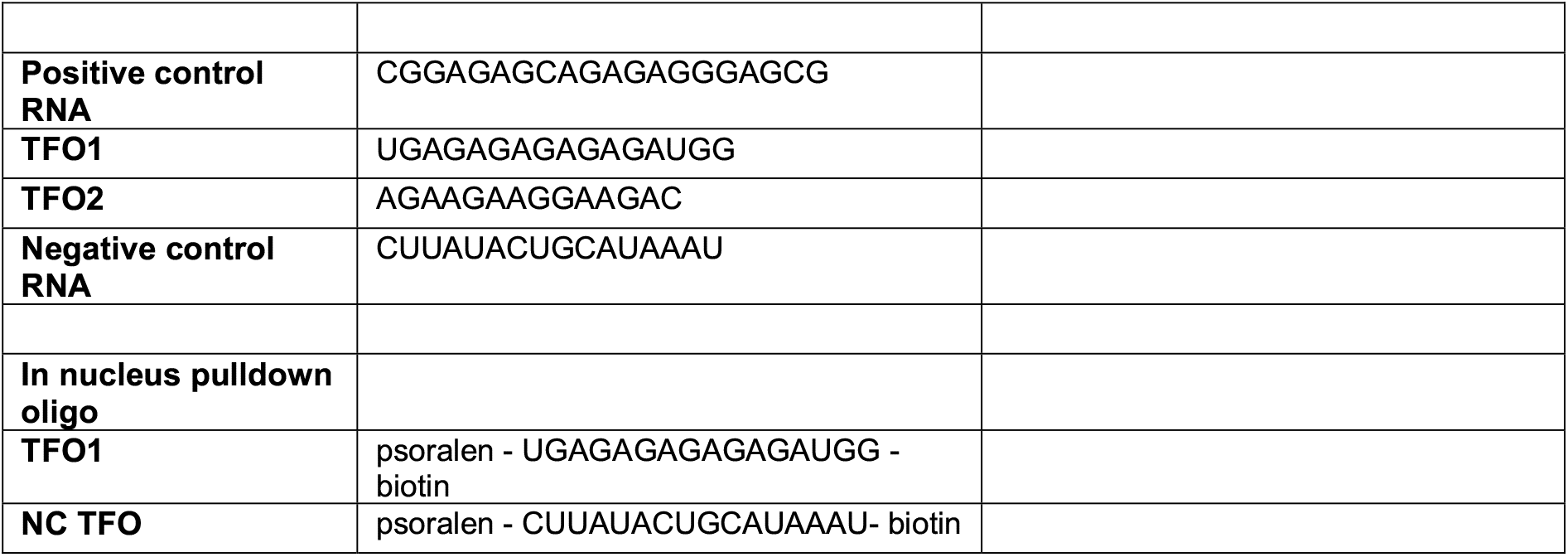
List of all primers and oligonucleotides used for the current study

**Fig. 2 supplementary:**
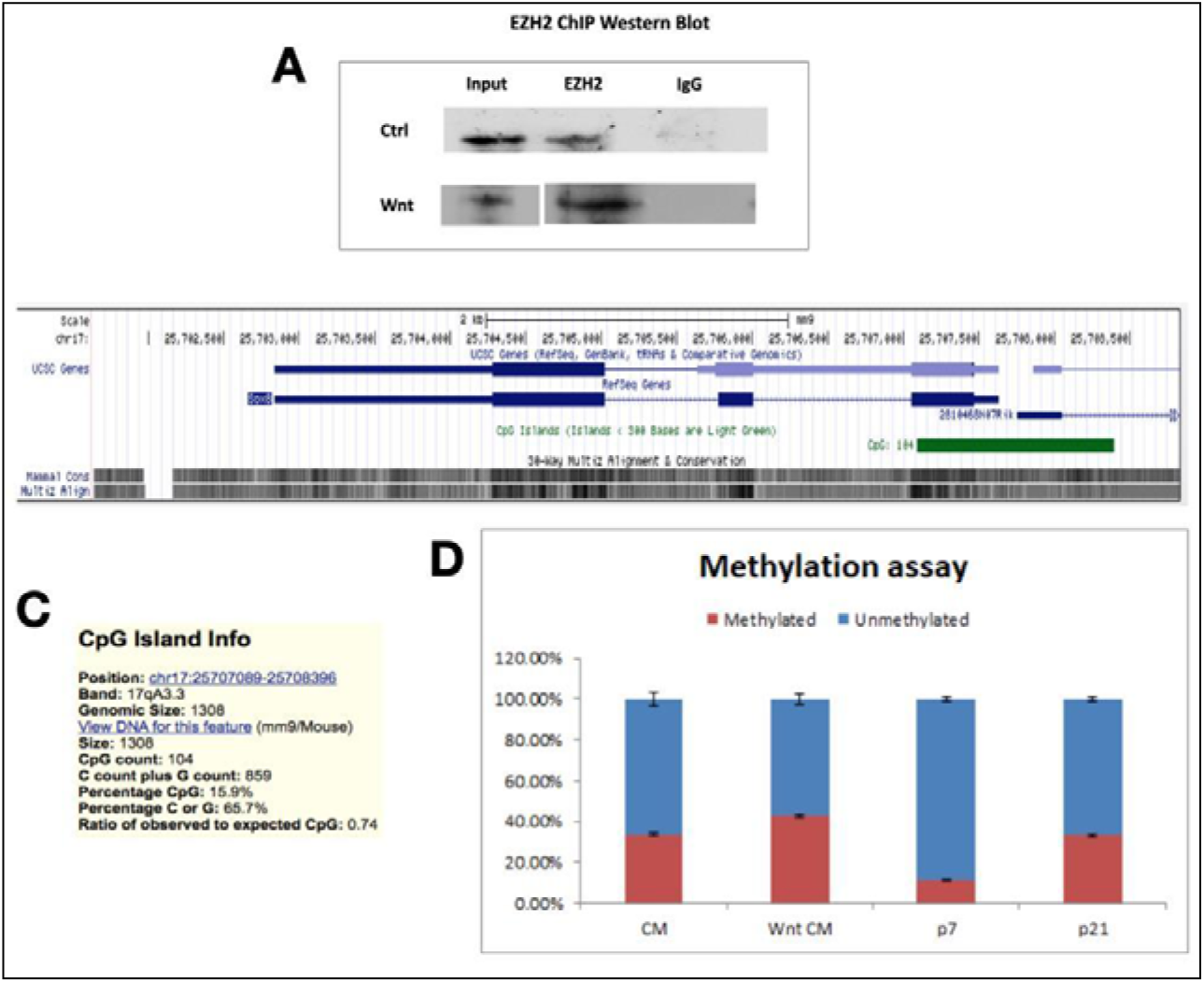
**(A)** ChIP wb for ezh2 performed in gcl-spg cells grown either in control or wnt media **(B)** CpG island encompassing the sox8 promoter **(C)** information about the CpG island **(D)** % CpG methylation at the sox8 cpg island in both cells (increase in wnt when compare to control) and mice testis (increase in p21 when compared to p7). Data in the graph has been plotted as mean ± s.d., n=3.

**Fig. 4 supplementary:**
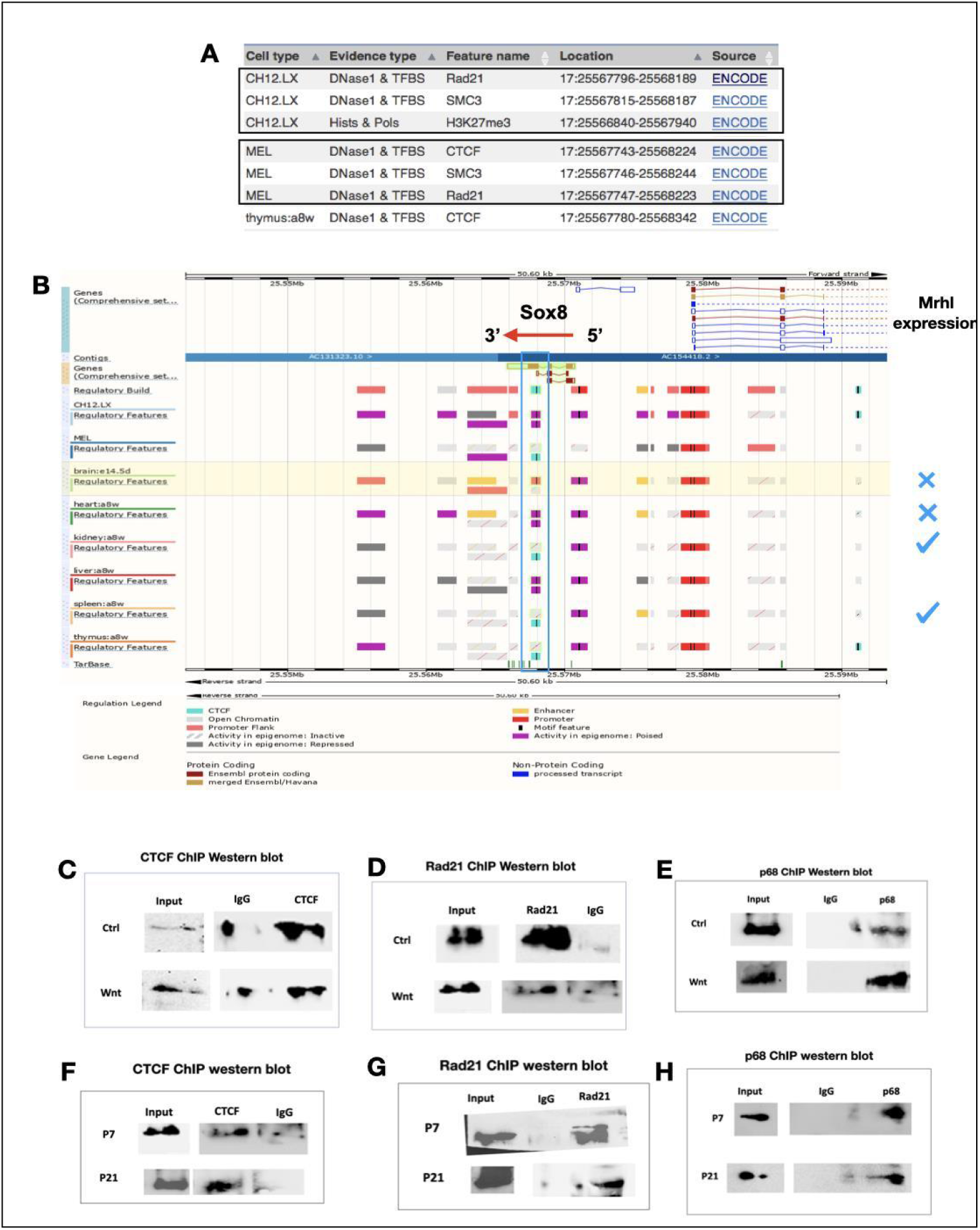
**(A)** Cohesin subunit SMC3 and Rad21 are found to bind close to the CTCF binding site in exon 3 of Sox8 **(B)** CTCF appears to be bound at the binding site within Sox8 in those tissue in which mrhl is expressed but not in some other in which mrhl isn’t expressed **(C)** ChIP WB for CTCF in Gcl-spg cells with and without Wnt induction. **(D)** ChIP WB for Rad21 in Gcl-spg cells with and without Wnt induction **(E)** ChIP WB for p68 in Gcl-spg cells with and without Wnt induction **(F)** ChIP WB for CTCF in mice testes of ages 7 days and 21 days **(G)** ChIP WB for Rad21 in mice testes of ages 7 days and 21 days **(H)** ChIP WB for p68 in mice testes of ages 7 days and 21 days

**Supplementary figure 5:**
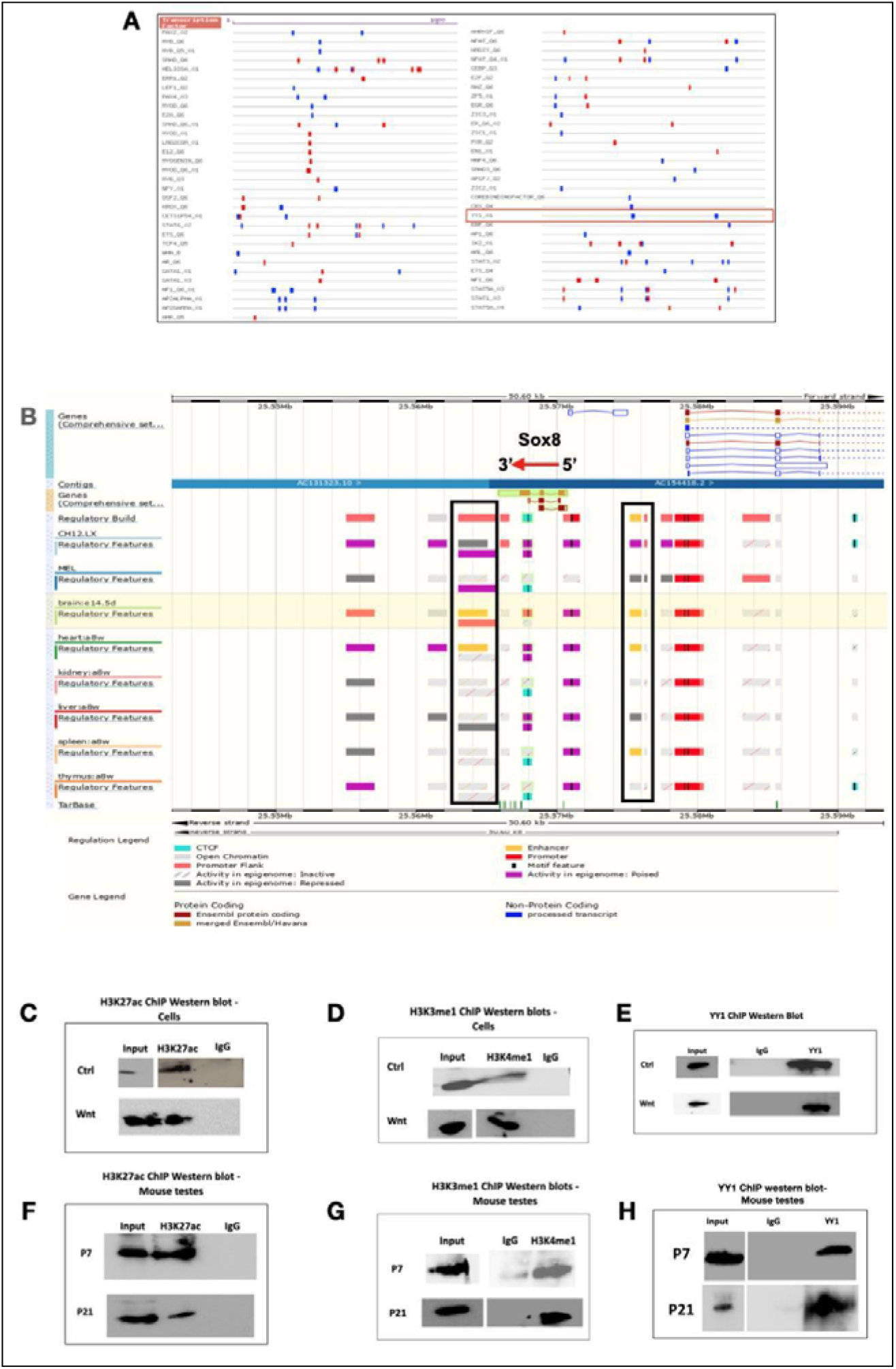
**(A)** Results from the Gene promoter miner tool to scan the promoter of Sox8 for presence of binding sites for transcription factors. YY1 has two binding sites in the promoter (highlighted with the red box). **(B)** The enhancer element present upstream and downstream of Sox8 (highlighted with black boxes) show activity corresponding to transcriptional activity of the Sox8 gene. **(C)** ChIP western blot for H3K27ac in Gcl-spg cells with and without Wnt activation **(D)** ChIP western blot for H3K4mel in Gcl-spg cells with and without Wnt activation **(E)** ChIP western blot for YY1 in Gclspg cells with and without Wnt activation **(F)** ChIP western blot for H3K27ac in P7 and P21 mice testes **(G)** ChIP western blot for H3K4mel in P7 and P21 mice testes **(H)** ChIP western blot for YY1 in P7 and P21 mice testes

**Supplementary figure 6:**
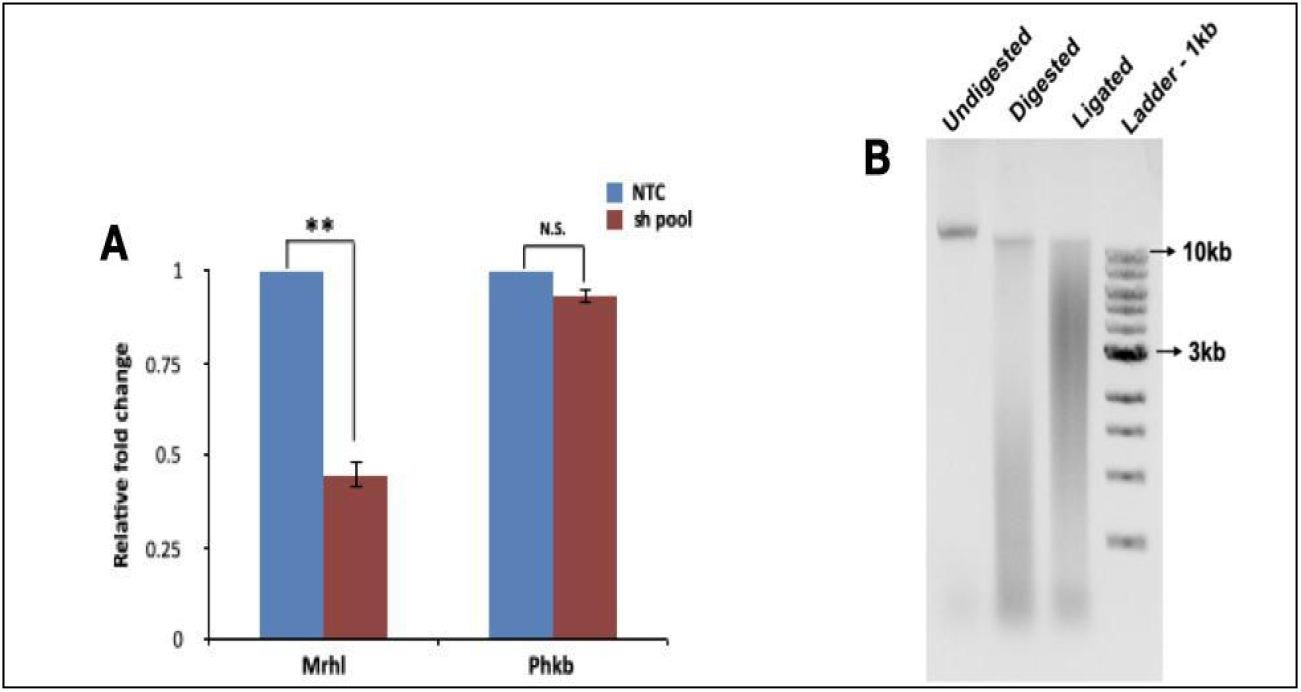
**(A)** To knockdown mrhl, a pool of both shRNA were used and silencing of around 55% was obtained while the host phkb gene expression level was not significantly perturbed. **(B)** Agarose gel image for DNA isolated from key steps of 3C shows enrichment of DNA below 3kb post restriction digestion with DpnII and an upward shift of the smear upon ligation

**Supplementary figure 7:**
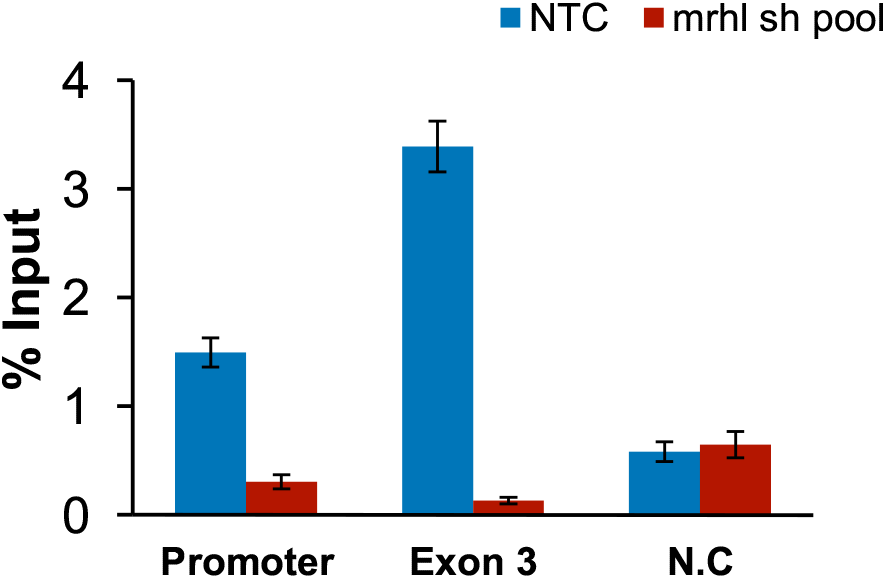
ChIP qPCR with CTCF specific antibody (no cross reactivity with CTCFL) shows the presence of CTCF at the Sox8 locus in the presence of mrhl with a reduction in its occupancy upon mrhl downregulation. Graph has been plotted as mean. N=3.

## References

1. Akhade, V. S., Arun, G., Donakonda, S., & Rao, M. R. (2014). Genome wide chromatin occupancy of mrhl RNA and its role in gene regulation in mouse spermatogonial cells. RNA Biol, 11(10), 1262–1279.

2. Akhade, V. S., Dighe, S. N., Kataruka, S., & Rao, M. R. (2016). Mechanism of Wnt signaling induced down regulation of mrhl long non-coding RNA in mouse spermatogonial cells. Nucleic Acids Res, 44(1), 387–401.

3. Barrionuevo, F., & Scherer, G. (2010). SOX E genes: SOX9 and SOX8 in mammalian testis development. IntJBiochem Cell Biol, 42(3), 433–436.

4. Barrionuevo, F. J., Hurtado, A., Kim, G. J., Real, F. M., Bakkali, M., Kopp, J. L., et al. (2016). Sox9 and Sox8 protect the adult testis from male-to-female genetic reprogramming and complete degeneration. Elife, 5.

5. Blank-Giwojna, A., Postepska-Igielska, A., & Grummt, I. (2019). lncRNA KHPS1 Activates a Poised Enhancer by Triplex-Dependent Recruitment of Epigenomic Regulators. Cell Rep, 26(11), 2904–2915.e2904.

6. Buske, F. A., Bauer, D. C., Mattick, J. S., & Bailey, T. L. (2012). Triplexator: detecting nucleic acid triple helices in genomic and transcriptomic data. Genome Res, 22(7), 1372–1381.

7. Essien, K., Vigneau, S., Apreleva, S., Singh, L. N., Bartolomei, M. S., & Hannenhalli, S. (2009). CTCF binding site classes exhibit distinct evolutionary, genomic, epigenomic and transcriptomic features. Genome Biol, 10(11), R131.

8. Feng, J., Liu, T., Qin, B., Zhang, Y., & Liu, X. S. (2012). Identifying ChIP-seq enrichment using MACS. NatProtoc, 7(9), 1728–1740.

9. Garcia-Moreno, S. A., Futtner, C. R., Salamone, I. M., Gonen, N., Lovell-Badge, R., & Maatouk, D. M. (2019). Gonadal supporting cells acquire sex-specific chromatin landscapes during mammalian sex determination. Dev Biol, 446(2), 168–179.

10. Gonen, N., Futtner, C. R., Wood, S., Garcia-Moreno, S. A., Salamone, I. M., Samson, S. C., et al. (2018). Sex reversal following deletion of a single distal enhancer of. Science, 360(6396), 1469–1473.

11. Grote, P., & Herrmann, B. G. (2013). The long non-coding RNA Fendrr links epigenetic control mechanisms to gene regulatory networks in mammalian embryogenesis. RNA Biol, 10(10), 1579–1585.

12. Guth, S. I., Bösl, M. R., Sock, E., & Wegner, M. (2010). Evolutionary conserved sequence elements with embryonic enhancer activity in the vicinity of the mammalian Sox8 gene. Int J Biochem Cell Biol, 42(3), 465–471.

13. Huang, D., Petrykowska, H. M., Miller, B. F., Elnitski, L., & Ovcharenko, I. (2019). Identification of human silencers by correlating cross-tissue epigenetic profiles and gene expression. Genome Res, 29(4), 657–667.

14. Kalwa, M., Hänzelmann, S., Otto, S., Kuo, C. C., Franzen, J., Joussen, S., et al. (2016). The lncRNA HOTAIR impacts on mesenchymal stem cells via triple helix formation. Nucleic Acids Res, 44(22), 10631–10643.

15. Kataruka, S., Akhade, V. S., Kayyar, B., & Rao, M. R. S. (2017). Mrhl Long Noncoding RNA Mediates Meiotic Commitment of Mouse Spermatogonial Cells by Regulating Sox8 Expression. Mol Cell Biol, 37(14).

16. Kentepozidou, E., Aitken, S. J., Feig, C., Stefflova, K., Ibarra-Soria, X., Odom, D. T., et al. (2020). Clustered CTCF binding is an evolutionary mechanism to maintain topologically associating domains. Genome Biol, 21(1), 5.

17. Kim, D., Pertea, G., Trapnell, C., Pimentel, H., Kelley, R., & Salzberg, S. L. (2013). TopHat2: accurate alignment of transcriptomes in the presence of insertions, deletions and gene fusions. Genome Biol, 14(4), R36.

18. Langmead, B., & Salzberg, S. L. (2012). Fast gapped-read alignment with Bowtie 2. Nat Methods, 9(4), 357–359.

19. Li, H., Handsaker, B., Wysoker, A., Fennell, T., Ruan, J., Homer, N., et al. (2009). The Sequence Alignment/Map format and SAMtools. Bioinformatics, 25(16), 2078–2079.

20. Mondal, T., Subhash, S., Vaid, R., Enroth, S., Uday, S., Reinius, B., et al. (2015). MEG3 long noncoding RNA regulates the TGF-β pathway genes through formation of RNA-DNA triplex structures. Nat Commun, 6, 7743.

21. Mumbach, M. R., Rubin, A. J., Flynn, R. A., Dai, C., Khavari, P. A., Greenleaf, W. J., et al. (2016). HiChIP: efficient and sensitive analysis of protein-directed genome architecture. Nat Methods, 73(11), 919–922.

22. Naumova, N., Smith, E. M., Zhan, Y., & Dekker, J. (2012). Analysis of long-range chromatin interactions using Chromosome Conformation Capture. Methods, 58(3), 192–203.

23. Nishant, K. T., Ravishankar, H., & Rao, M. R. (2004). Characterization of a mouse recombination hot spot locus encoding a novel non-protein-coding RNA. Mol Cell Biol, 24(12), 5620–5634.

24. O’Bryan, M. K., Takada, S., Kennedy, C. L., Scott, G., Harada, S., Ray, M. K., et al. (2008). Sox8 is a critical regulator of adult Sertoli cell function and male fertility. Dev Biol, 316(2), 359–370.

25. O’Leary, V. B., Ovsepian, S. V., Carrascosa, L. G., Buske, F. A., Radulovic, V., Niyazi, M., et al. (2015). PARTICLE, a Triplex-Forming Long ncRNA, Regulates Locus-Specific Methylation in Response to Low-Dose Irradiation. Cell Rep, 77(3), 474–485.

26. Ogboume, S., & Antalis, T. M. (1998). Transcriptional control and the role of silencers in transcriptional regulation in eukaryotes. Biochem J, 331 (Pt 1), 1–14.

27. Ong, C. T., & Corces, V. G. (2014). CTCF: an architectural protein bridging genome topology and function. Nat Rev Genet, 15(4), 234–246.

28. Pal, D., Neha, C. V., Bhaduri, U., Zenia, Z., Dutta, S., Chidambaram, S., et al. (2021). LncRNA Mrhl orchestrates differentiation programs in mouse embryonic stem cells through chromatin mediated regulation. Stem Cell Res, 53, 102250.

29. Pentland, I., Campos-León, K., Cotic, M., Davies, K. J., Wood, C. D., Groves, I. J., et al. (2018). Disruption of CTCF-YY1-dependent looping of the human papillomavirus genome activates differentiation-induced viral oncogene transcription. PLoSBiol, 16(10), e2005752.

30. Postepska-Igielska, A., Giwojna, A., Gasri-Plotnitsky, L., Schmitt, N., Dold, A., Ginsberg, D., et al. (2015). LncRNA Khpsl Regulates Expression of the Proto-oncogene SPHK1 via Triplex-Mediated Changes in Chromatin Structure. Mol Cell, 60(4), 626–636.

31. Richardson, N., Gillot, I., Gregoire, E. P., Youssef, S. A., de Rooij, D., de Bruin, A., et al. (2020). and. Elife, 9.

32. Robinson, J. T., Thorvaldsdóttir, H., Winckler, W., Guttman, M., Lander, E. S., Getz, G., et al. (2011). Integrative genomics viewer. Nat Biotechnol, 29(1), 24–26.

33. Saldaña-Meyer, R., González-Buendía, E., Guerrero, G., Narendra, V., Bonasio, R., Recillas-Targa, F., et al. (2014). CTCF regulates the human p53 gene through direct interaction with its natural antisense transcript, Wrap53. Genes Dev, 28(7), 723–734.

34. Saldaña-Meyer, R., Rodriguez-Hemaez, J., Escobar, T., Nishana, M., Jácome-López, K., Nora, E. P., et al. (2019). RNA Interactions Are Essential for CTCF-Mediated Genome Organization. Mol Cell, 76(3), 412–422.e415.

35. Schmitz, K. M., Mayer, C., Postepska, A., & Grummt, I. (2010). Interaction of noncoding RNA with the rDNA promoter mediates recruitment of DNMT3b and silencing of rRNA genes. Genes Dev, 24(20), 2264–2269.

36. Singh, A. P., Harada, S., & Mishina, Y. (2009). Downstream genes of Sox8 that would affect adult male fertility. Sex Dev, 3(1), 16–25.

37. Trapnell, C., Williams, B. A., Pertea, G., Mortazavi, A., Kwan, G., van Baren, M. J., et al. (2010). Transcript assembly and quantification by RNA-Seq reveals unannotated transcripts and isoform switching during cell differentiation. Nat Biotechnol, 28(5), 511–515.

38. Weintraub, A. S., Li, C. H., Zamudio, A. V., Sigova, A. A., Hannett, N. M., Day, D. S., et al. (2017). YY1 Is a Structural Regulator of Enhancer-Promoter Loops. Cell, 171(7), 1573–1588.el528.

39. Xiang, J. F., Yin, Q. F., Chen, T., Zhang, Y., Zhang, X. O., Wu, Z., et al. (2014). Human colorectal cancer-specific CCAT1-L lncRNA regulates long-range chromatin interactions at the MYC locus. Cell Res, 24(5), 513–531.

40. Yang, Y., & Li, G. (2020). Post-translational modifications of PRC2: signals directing its activity. Epigenetics Chromatin, 13(1), 47.

41. Yao, H., Brick, K., Evrard, Y., Xiao, T., Camerini-Otero, R. D., & Felsenfeld, G. (2010). Mediation of CTCF transcriptional insulation by DEAD-box RNA-binding protein p68 and steroid receptor RNA activator SRA. Genes Dev, 24(22), 2543–2555.

42. Yao H, Brick K, Evrard Y, Xiao T, Camerini-Otero RD, Felsenfeld G. 2010. Mediation of CTCF transcriptional insulation by DEAD-box RNA-binding protein p68 and steroid receptor RNA activator SRA. Genes Dev 24:2543–2555.

